# Profiling Peripheral Blood with an Optimized, Multiplexed, Single-cell Multiome Approach Supports an Insulin-driven Asthma Subtype

**DOI:** 10.64898/2026.03.27.714744

**Authors:** Jiacheng Ding, HyunJin Kang, Amber L. Spangenberg, Yusha Liu, Fernando D. Martinez, Tara F. Carr, Darren A. Cusanovich

**Author notes:** Corresponding Authors: Tara F. Carr and Darren A. Cusanovich.

## Abstract

RNA sequencing (RNA-seq) and the Assay for Transposase-Accessible Chromatin using sequencing (ATAC-seq) have become standard techniques for studying gene regulation in human populations. Single-cell (sc) “multiomic” genomic methodologies now enable researchers to dissect cellular heterogeneity while simultaneously measuring gene expression and chromatin accessibility within individual cells. However, single-cell approaches remain experimentally complex and cost-prohibitive, limiting their application in population studies, and motivating the development of new strategies for population-scale single-cell investigations. To this end, we have adapted and optimized a previous multiomic protocol, “Transcriptome, Epitope, and ATAC sequencing” (TEA-seq) through experimentation and simulation to incorporate sample multiplexing, thus resulting in our “multiplexed TEA-seq” (mTEA-seq) protocol. Using mTEA-seq, we sought to determine whether asthma that develops in conjunction with early-life elevated insulin levels might have an identifiable molecular signature. We studied samples from adult individuals (54 subjects, 272,003 cells) from the Tucson Children’s Respiratory Study (TCRS), a birth cohort phenotypically characterized over four decades, to identify unique molecular characteristics of blood cells from asthmatics who had high serum insulin levels at age 6. Using a Bayesian approach, we found striking sex-specific effects. Male asthmatic subjects with high insulin at age 6 displayed widespread immune transcriptional and epigenetic alterations into adulthood compared to male non-asthmatic subjects without elevated insulin at age 6. We also found that male non-asthmatics with early-life high insulin showed epigenetic perturbations in adulthood, but not transcriptional changes. The consistency of epigenetic signals between these two groups that had high insulin at age 6 was highly cell-type-specific. For example, CD14+ monocytes displayed broadly common insulin-associated chromatin remodeling regardless of asthma status, while NK cells exhibited unique patterns of insulin-associated epigenetic reprogramming depending on asthma status. Finally, genotyping performed directly from our single-cell data enabled cell type-specific cis-QTL mapping that suggested HLA-DQB1 and AHI as genes for future study in insulin-associated asthma. Our investigation of childhood insulin-associated asthma demonstrates a metabolically-driven alterations on immune cells persisting into adulthood, thus providing a molecular signature of this asthma subtype, and offering novel insights for disease prevention and therapeutic intervention.

## Introduction

Since the development of single-cell sequencing technologies in 2009^1,2^, they have emerged as transformative tools, enabling high-resolution profiling across various molecular modalities, including transcriptomics^3–6^, DNA methylation^7,8^, chromatin accessibility^9–13^, chromatin conformation^14^, and proteomics^15,16^. These innovations have considerably advanced research in complex diseases, developmental biology, and oncology by providing deeper insights into cellular heterogeneity. A second wave of single-cell genomic technology development has resulted in techniques for the simultaneous measurement of multiple molecular layers within individual cells, known as single-cell multiomics or co-assay approaches^17–21^ (e.g., parallel profiling of the transcriptome and epigenome). Despite these technological advances, challenges remain. The inherently low abundance of target molecules (e.g., mRNA) and suboptimal capture efficiencies in individual cells^22^ result in data characterized by excessive zero inflation. Such dropout events complicate statistical analyses, as models may misinterpret zeros as true biological absence and overestimate differences among low-expression features, thereby inflating false-positive rates^23^. These issues are further compounded by the frequent use of few or no biological replicates in single-cell studies, often driven by cost concerns, which can lead to inflated false-positive rates among highly expressed features as well^23^.

Although combinatorial indexing and related technologies have substantially improved sample multiplexing capacity and throughput, these methods often require labor-intensive, time-consuming workflows and frequently involve sample fixation to preserve molecular integrity across multiple rounds of barcoding, all of which can span several days. Consequently, there remains a strong demand for single-cell multiomic platforms that are user-friendly, time- and labor-efficient, cost-effective, and capable of generating large-scale, high-quality datasets. In response to this need, we have expanded the “Transcriptome, Epitope, and ATAC sequencing” (TEA-seq) method^21^ to allow for sample multiplexing, which we call “mTEA-seq”. mTEA-seq is a single-cell multiomic approach that integrates cell hashing, the 10X Genomics Multiome assay, and several other important optimizations specific to peripheral blood mononuclear cells (PBMCs). As we show, this method enables rapid, reliable, and scalable profiling of both the transcriptome and epigenome while maintaining cost-efficiency, facilitating broader applications in large-cohort studies.

As a test case for “multiplexed TEA-seq” (mTEA-seq), we sought to explore the immune cell state of subjects in a longitudinal cohort that was established to characterize the dynamics of asthma pathogenesis, the Tucson Children’s Respiratory Study (TCRS)^24^. Asthma is a complex disease (i.e., shaped by both genetic and environmental influences^25^) with documented phenotypic and endotypic heterogeneity^26^ and pronounced sexual dimorphism^27–30^. The molecular pathways mediating these differences are only beginning to be explored. Recent single-cell transcriptomic studies have begun elucidating immune cell landscapes in the human lung^31^, bronchoalveolar lavage^32^, sputum^33^, and PBMCs^34,35^, however, integrated single-cell transcriptomic and epigenomic analyses remain limited, hindering a comprehensive understanding of how functional genomic modules interact with disease-associated genes. Moreover, the high costs and sample access limitations have constrained most single-cell asthma studies to relatively small numbers of subjects, thereby reducing their statistical power.

While no single study can address all of these gaps, TCRS provides a valuable resource for investigating early-life asthma origins. As the longest ongoing birth cohort focused on asthma and airway diseases, TCRS offers the advantages of extensive longitudinal data collection^24^. A recent TCRS report^36^ identified an association between elevated serum insulin levels measured at age six and an increased subsequent prevalence of active asthma from childhood through adulthood, independent of body mass index. Importantly, this finding was replicated in a second cohort, the Avon Longitudinal Study of Parents and Children (ALSPAC)^37–39^. Although obesity, metabolic dysfunction, and insulin resistance have been implicated in asthma pathogenesis (e.g., ^40–43^), elevated serum insulin in early life appears to represent an independent risk factor, thus potentially defining a distinct asthma endotype. Given that insulin has cell-type-specific pleiotropic influences on cellular function, including in the immune system (reviewed in ^44^), the observed link between insulin and asthma raised intriguing questions about whether there may be a molecular signature that would define such an asthma endotype persisting into adulthood. To investigate this possibility, we isolated PBMCs from 54 current participants in the TCRS cohort (∼40 years old), for whom serum insulin levels had been measured at age six. Participants were stratified into four phenotypic groups based on early-life insulin levels and asthma ever diagnosis, samples were profiled via mTEA-seq, and we investigated the molecular signatures of this potential insulin-associated asthma endotype with state-of-the-art approaches.

## Results

### Optimizing a multiplexed multiomic assay

Given the necessity of biological replication for single-cell omic profiling and the challenges (both economic and logistical) of conducting single-cell studies incorporating many samples, we sought to establish a working protocol for multiplex processing of PBMC samples adapted from the previously published TEA-seq assay^21^. To do so, we optimized a multiplex processing protocol for PBMC samples utilizing antibody-based cellular hashing for unambiguous sample multiplexing, FACS depletion of dead cells and granulocytes, and multiomic processing on the 10X Genomics platform (**Fig. 1A**). Key protocol optimizations included adding CD16 antibody to the staining cocktail alongside CD15 antibody to better distinguish monocytes from neutrophils during FACS sorting (**Fig. S1**) and conducting independent amplification reactions for cDNA and hashtag oligos (HTOs) rather than co-amplifying them, which markedly reduced non-specific byproducts and PCR bubble artifacts while simplifying the protocol (**Fig. S2**). Through simulations to model cell singlet and multiplet distributions for microfluidic systems with cell inputs ranging from 0 to 200,000, we determined that optimal cell loading at approximately 48,484 cells would maximize the yield of singlets over multiplets, generating 20,584 singlet droplets (**Fig. 1B**). In an initial pilot experiment, we multiplexed PBMCs from 4 individuals (loading 40,000 cells and recovering 16,675 singlets) and found the results to meet our expectations of quality, with all individuals well represented across the dataset (**Fig. 1C**, **Fig. S3**, See Supplementary Note for more details on optimization of the protocol).

**Figure 1.**
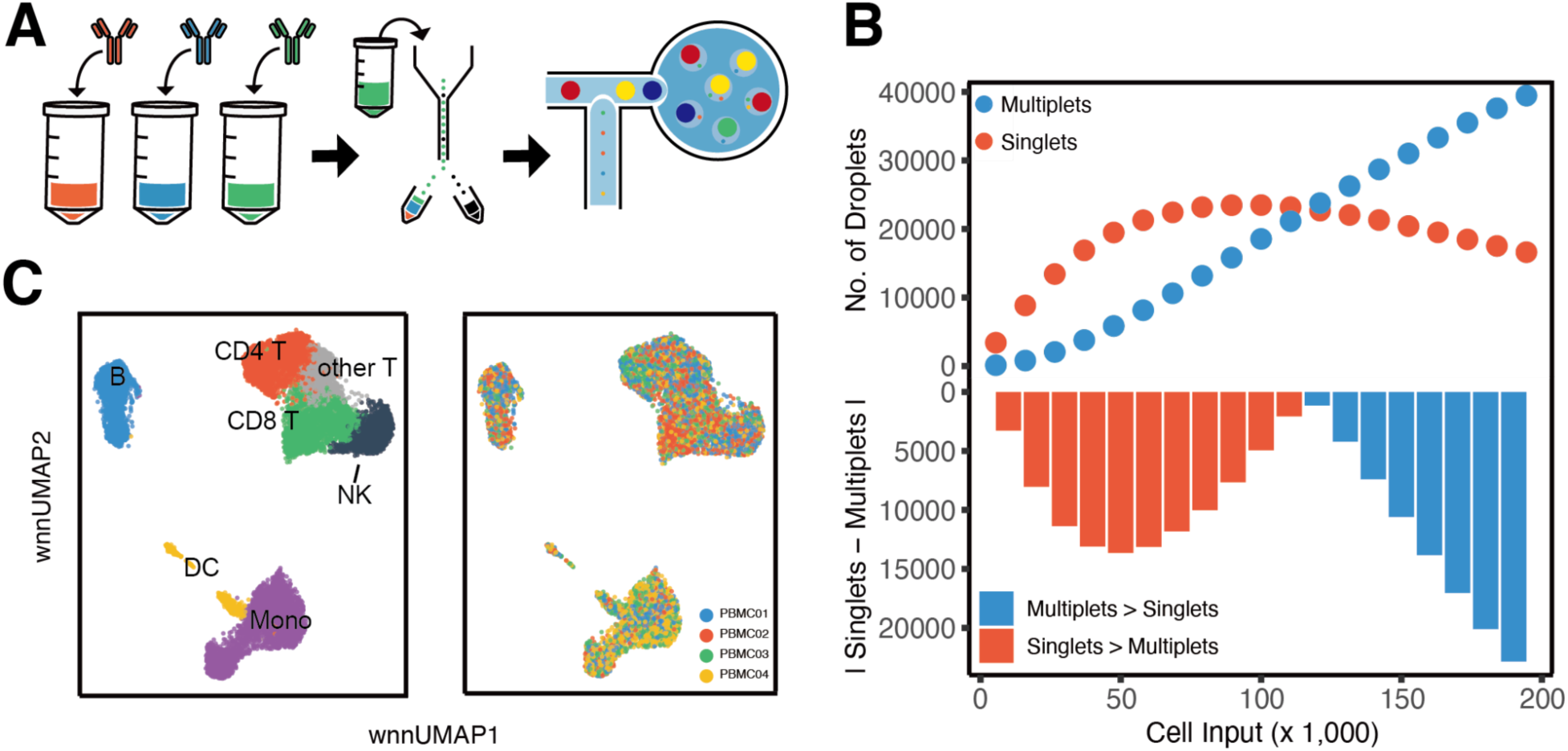
mTEA-seq method validation. **(A)** Schematic overview of mTEA-seq. PBMC samples are stained with an antibody cocktail in addition to hashing antibodies, then FACS sorted into one tube. Pooled cells are ultimately processed through 10X multiomic assay. **(B)** Simulation of singlet and multiplet rates with cell input ranging from 0 to 200,000. Upper panel: a dot plot showing how the number of singlet droplets (red) and multiplet droplets (blue) vary with different cell inputs. Lower panel: a bar plot showing the absolute difference in number of singlets and multiplets. Bar colors indicate whether there are more singlets (red) or multiplets (blue). **(C)** Weighted nearest neighbor (wnn) UMAP colored by predicted cell types (left) and subjects (right).

### Profiling PBMCs from individuals stratified by early-life insulin measurements and asthma status

Having established that mTEA-seq was capable of producing high-quality single-cell multimodal data in multiplexed samples, we next sought to apply our method to TCRS samples^36^. To determine the immunological landscape of individuals with early-life elevated insulin and assess if those with high insulin and an asthma diagnosis might represent a distinct asthma endotype, we profiled PBMCs collected at 40 years old from TCRS participants stratified by early-life insulin level (high vs low) and asthma status (ever diagnosed vs never diagnosed) using mTEA-seq. This resulted in four groups: low insulin non-asthmatics (LN), low insulin asthmatics (LA), high insulin non-asthmatics (HN), and high insulin asthmatics (HA).

After excluding low viability samples (**Fig. S4**), we generated high-quality single-cell multiome data on 272,003 PBMCs representing 54 unique individuals. Data were integrated, clustered, and used to produce a unified UMAP based on both RNA and ATAC data (**Fig. 2A**). Using a previously annotated PBMC reference^45^, we identified 19 cell types. Across the study, the relative proportions of these cell types were in keeping with reference ranges for PBMCs^46,47^. To confirm cell type annotations, cell type markers for each cell type were compared with the reference PBMC data (**Fig. S5**). Our initial exploratory analyses suggested that there were considerable differences in the profiles of male and female subjects (**Fig. S6**), and given the known sex-based differences in asthma prevalence^48^, insulin molecular pathways^49^, and insulin resistance prevalence^50^, we decided to proceed by considering the two sexes separately for all subsequent analyses.

**Figure 2.**
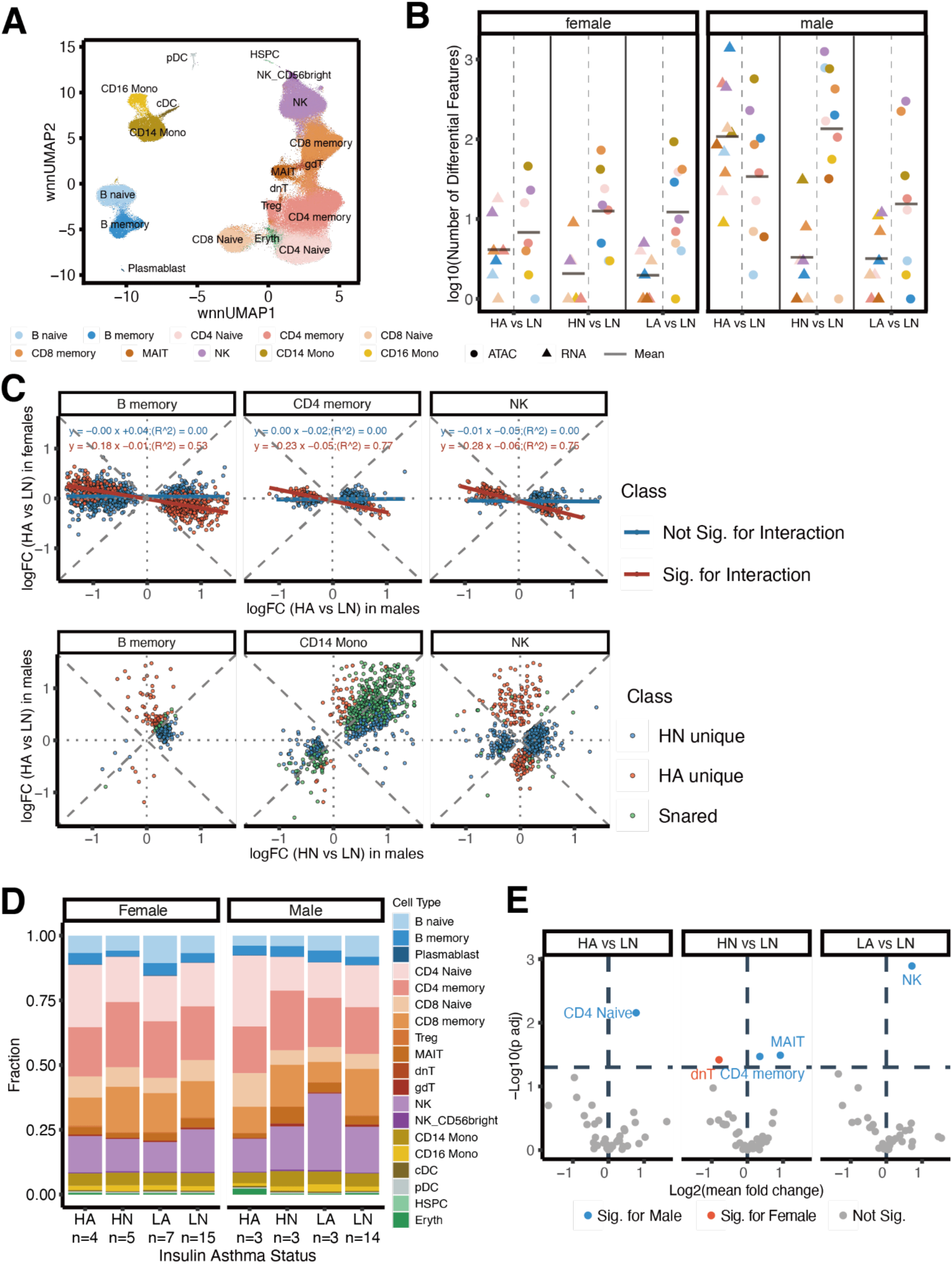
Cell type-specific differential analysis with TCRS PBMC single-cell multiomic data. **(A)** wnn UMAP plot of 272,003 PBMCs from 75 samples, representing 54 distinct individuals, colored by cell type. **(B)** Strip plot depicting the count of significantly altered molecular features per cell type and assay, stratified by sex. Triangles represent differentially expressed genes, while circles denote differentially accessible chromatin regions. Color indicates cell type (female: left panel; male: right panel). **(C)** Scatter plot displaying log2(fold-change) comparison by cell type between groups. Top panel showing DE log2(fold-change) comparisons between female and male HA vs LN tests. Color codes for genes categorized as not significant for a sex interaction (red) and significant for an interaction (blue). Lines indicated regression curves. Bottom panel showing DA log2(fold-change) comparisons between male HA vs LN and HN vs LN tests. Color codes for chromatin regions that are shared (green), unique for HN (blue), or unique for HA (red). **(D)** Stacked bar plot illustrating the average proportion of annotated cell types across groups categorized by insulin and asthma status - low insulin non-asthmatic (LN), low insulin asthmatic (LA), high insulin non-asthmatic (HN), and high insulin asthmatic (HA) - further stratified by sex (female: left panel; male: right panel). **(E)** Volcano plots illustrate the outcomes of differential analyses in cell type proportions, based on t-tests comparing the "HA," "HN," and LA groups against the LN reference group within sex-stratified populations. The x-axis represents the log_2_-transformed mean fold-change, while the y-axis indicates the -log_10_(adj. p-values). Dashed lines denote the threshold for statistical significance at an adj. p-value of 0.05. Data points are color-coded according to significance and sex specificity: non-significant (grey), significant in males (blue), and significant in females (red).

### Multivariate adaptive shrinkage identifies sex- and condition-specific molecular signatures

After annotating cell types, we aggregated single-cell gene expression and chromatin accessibility data by individual and cell type to perform “pseudo-bulk” differential tests^23^. To carry out these tests, we chose to adopt a recently developed Bayesian framework, called Multivariate Adaptive Shrinkage (mash), implemented in the mashr package in R^51^, which allowed us to jointly model all of the comparisons in a given cell type in order to increase power, while focusing our analyses on how the case groups compared to the controls (i.e., HA vs LN, HN vs LN, and LA vs LN), and being able to control for covariates such as repeated measures (see Methods, **Fig. S7**).

In total, we identified 2,974 differentially expressed (DE) genes across these comparisons at a 5% local false sign rate (lfsr). Of these, 2,895 DE genes were identified in males, while only 79 were identified in females, revealing a striking sex-specific effect. Notably, the majority of significant gene expression differences in males were greatly concentrated in the HA vs LN comparison, where 2,807 DE genes were identified in this comparison, in contrast to 49 DE genes for HN vs LN and 39 DE genes for LA vs LN (**Fig. 2B**). Among the cell types with significant differentially expressed genes in the male HA vs LN comparison, memory B cells stood out as the cell type with the most significant DE genes (N = 1,385), followed by CD4+ memory T cells and NK cells, with 496 and 448 DE genes, respectively. By way of contrast, the most DE genes for any comparison in females was 18 genes DE for CD4+ naive T cells in the HA vs LN comparison.

To further investigate whether there was statistical support for the transcriptional changes identified in the male HA vs LN comparison to be truly specific to males, we performed an interaction analysis incorporating both HA phenotype and sex in a linear model (see Methods: Evaluation of sex-specific transcription response using interaction modeling). This approach tested whether the expression differences observed in males were significantly different from those in females. Across all genes that were significantly differentially expressed in the male HA vs LN comparison, an average of 36.6% (median = 37%) also showed a significant sex interaction, indicating that over one-third of the transcriptional changes in this comparison were rigorously male-specific. To visualize this relationship, we compared the log₂ fold-changes (logFC) between males and females for the HA vs LN contrast across the three cell types that exhibited the strongest transcriptional (**Fig. 2C**): memory B cells, CD4 Memory cells, and NK cells. Genes with a significant sex interaction displayed a negative correlation between male and female logFC, consistent with the conservative nature of this test, whereas the genes without a significant interaction effect still showed little or no correlation, suggesting that the true proportion of differential genes with sex-specific effects is likely much larger than one third.

To identify regions of the genome with significantly differential chromatin accessibility (DA) between groups, we applied the same mashr pipeline (5% lfsr threshold) as was used for the transcriptomic analysis. As was the case for the RNA analysis, the majority of DA peaks were identified in males (5,449 for males and 505 for females). Interestingly, unlike the RNA analysis, which identified the HA vs LN comparison as the most transcriptionally different, the largest number of significant DA peaks was observed in the male HN vs LN comparison (N = 3,794), followed by the male HA vs LN comparison (N = 1,059). For example, although CD14+ monocytes exhibited a relatively small number of DE genes (N = 31) in the male HN vs LN comparison, they had a substantial number of DA chromatin regions (N =764). The major cell types exhibiting DA peaks in the male HA vs LN comparison were CD14⁺ monocytes (N=571), NK cells (N = 229), and memory B cells (N = 103). Among the three “case” group comparisons, the LA vs LN comparison demonstrated the lowest chromatin accessibility variation, with 596 total DA peaks identified in males and 230 in females.

Because both HN vs LN and HA vs LN exhibited so many DA sites, we wondered if these were correlated. To address this question, we took an approach similar to our analysis of sex-specific transcriptional responses, collecting all DA peaks in either test and comparing the LogFC between HN vs LN and HA vs LN for these sites. This analysis revealed varying degrees of overlap in chromatin regulation depending on the cell type. Among all cell types, CD14⁺ monocytes showed the greatest degree of concordance, with 513 shared significant peaks (62.4%) out of 822 total differential peaks, indicating a large set of regulatory elements jointly altered in both the HA and HN groups (**Fig. 2C**, lower panel). In contrast, memory B cells exhibited more distinct accessibility profiles, with only 12.6% of their significant DA peaks shared between the two comparisons. NK cells, despite displaying the largest total number of DA peaks (N = 1,453), showed only 2.1% overlap between HA and HN, suggesting largely condition-specific regulatory programs. These results highlight that CD14 monocytes may represent a cell type with shared chromatin response to insulin exposure, while B and NK cells display more specialized accessibility changes distinguishing the asthma-associated (HA) and non-asthmatic (HN) high insulin phenotypes.

Taken together, these results indicate that the HA group (i.e. individuals with early-life elevated serum insulin and asthma diagnoses) is both transcriptionally and epigenetically rewired relative to the LN group. On the other hand, the HN group (i.e. individuals with early-life serum insulin increases, but no asthma diagnosis) is dramatically epigenetically rewired, but does not exhibit a concomitant change in the transcriptional landscape. In contrast, the LA group (individuals diagnosed with asthma, but no early-life insulin elevation) displayed minimal changes at both the transcriptional and epigenetic levels.

To explore whether changes in gene expression or chromatin accessibility were accompanied by alterations in cellular composition, we also tested for significant differences in cell type proportions between the individuals in the three “case” groups (HA, HN, LA) and the LN controls (**Fig. 2D,E, Fig. S8**), after stratifying by sex. Overall, relatively few significant changes in cell type proportions were detected, indicating that most molecular alterations occurred without major shifts in cellular abundance. Notably, NK cells exhibited both a significant increase in proportion in the male LA group (adj. p-value = 1.28 × 10^-3^, logFC = 0.72) and a relatively large number of differentially accessible chromatin regions (N = 312). In contrast, other cell types with strong molecular signatures, such as memory B cells in the male HA group, did not show a significant change in abundance. Additional significant compositional changes included increased CD4⁺ naive T cells in the male HA group (adj. p-value = 6.97 × 10^-3^, logFC = 0.78), as well as increased mucosal-associated invariant T (MAIT) cells (adj. p-value = 3.24 × 10^-2^, logFC = 0.92) and CD4⁺ memory T cells (adj. p-value = 3.38 × 10^-2^, logFC = 0.35) in the male HN group. In females, double-negative T (dnT) cells were decreased in the HN group (adj. p-value = 3.82 × 10^-2^, logFC = -0.80).

### Pathway and motif analysis reveals metabolic, immune, and inflammatory reprogramming in high-insulin asthmatic males

To assess pathways implicated in our differential analysis, we performed over-representation analysis (ORA) using genes associated with either differential expression or differential accessibility (See Methods: Pathway over-representation analysis). We noted several pathways related to insulin and metabolism enriched in this analysis. For differential genes associated with the HA vs LN comparison (**Fig. 3A**), all five cell types tested were enriched for the term *‘positive regulation of metabolic process’* - memory B cells (adj. p-value = 3.83 × 10^-14^), CD14+ monocytes (adj. p-value =3.51 x 10^-2^), CD4+ memory T cells (adj. p-value = 4.00 × 10^-2^), CD8+ naive T cells (adj. p-value = 2.95 × 10^-2^), and NK cells (adj. p-value = 2.54 × 10^-4^). In addition, the *‘insulin resistance’* pathway was significant for memory B cells (adj. p-value = 8.23 × 10^-3^). Further, we identified significantly over-represented pathways related to insulin uptake, including *MAPK* (adj. p-value = 7.23 × 10^-6^), *FOXO* (adj. p-value = 7.75 × 10^-3^), and *NFkB* signaling (adj. p-value = 2.36 x 10^-2^) in memory B cells, all of which are well-studied pathways involved in regulating cell proliferation^52,53^, differentiation^54^, metabolism^55–57^, and immune response^53,57,58^. At the same time, we also identified many immune-relevant pathways enriched in this comparison. For example, *‘Th1 and Th2 cell differentiation’* was enriched in CD4+ memory T cells in the HA group (adj. p-value = 1.34 × 10^-2^), consistent with an imbalance in the ratio of Th1 and Th2 cells underlying the classic model of asthma pathogenesis^59^. Similarly, *‘Th17 cell differentiation’* was enriched in CD4+ memory T cells (adj. p-value = 5.59 × 10^-3^), consistent with the association of increased Th17+ T cells with severe asthma^60^.

**Figure 3.**
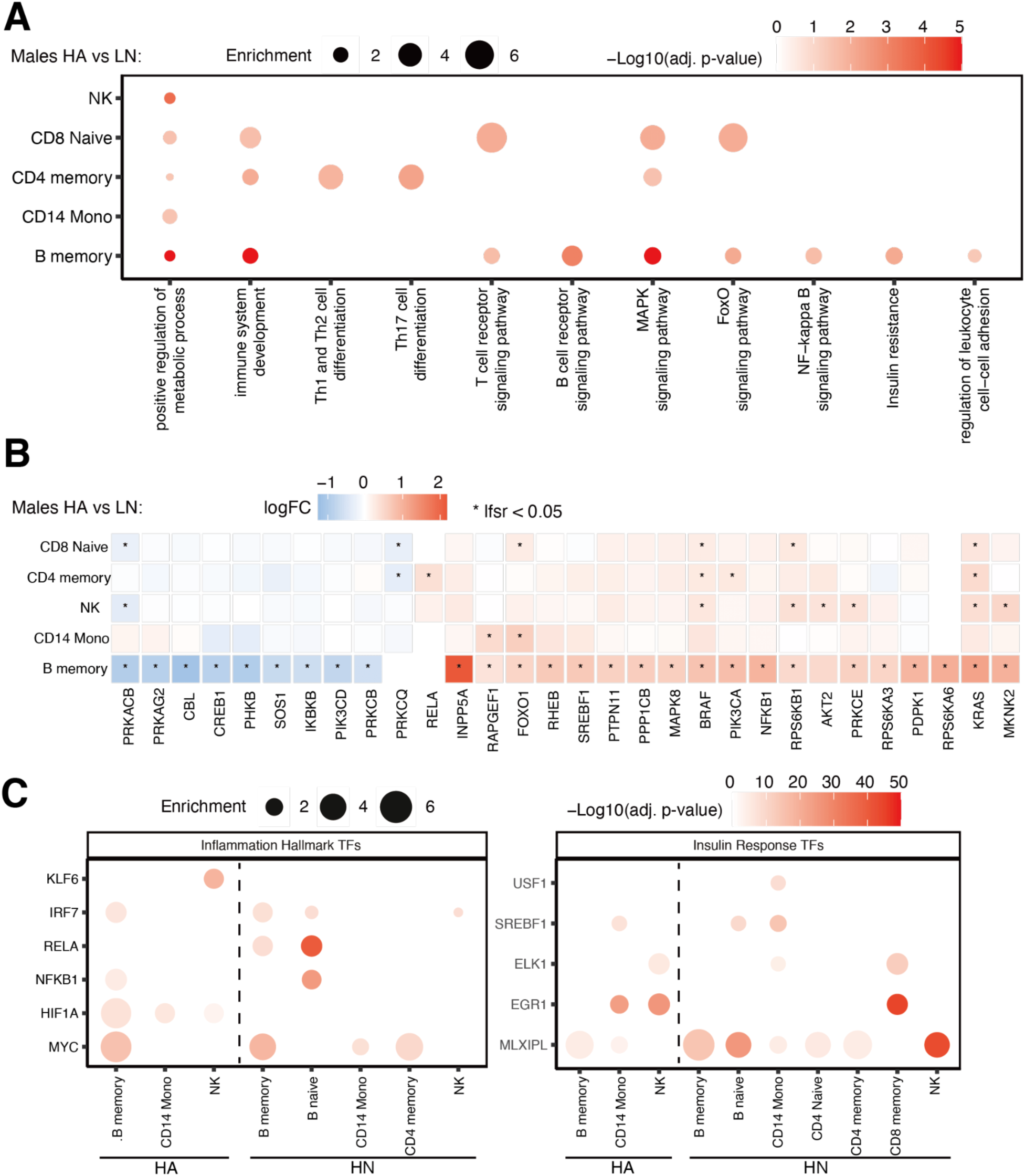
Cell type–specific pathway over-representation and differential expression of insulin signaling genes. **(A)** Dot plot summarizing ORA results for male HA vs LN comparison. Dot color represents statistical significance, expressed as the -log10(adjusted p-value), while dot size denotes the enrichment score. Only pathways meeting the significance criterion are included. **(B)** Heatmap displaying top genes involved in the insulin signaling pathway. Color reflects the absolute log2(fold-change) between case and control. An asterisk indicates statistical significance. **(C)** Dot plot summarizing motif enrichment analysis for cell types surpassing a predefined feature selection threshold. Plots were faceted into two TF panels. HA and HN data within each panel were segregated by a dashed line. Dot color represents statistical significance, expressed as the -log10(adjusted p-value), while dot size denotes the enrichment score.

Drilling down on the pathway-level enrichments (**Fig. 3A**), we next examined the expression of individual genes involved in the KEGG *‘insulin signaling’* and *‘insulin resistance’* pathways to pinpoint transcriptional alterations underlying the metabolic dysregulation observed in males. The top 30 insulin-related genes were selected based on a weighted score combining effect size and statistical significance across cell types. Multiple signaling components downstream of the insulin receptor were differentially expressed (**Fig. 3B**), predominantly in memory B cells. Among these were 4 different kinases involved in phosphoinositide metabolism, including Phosphatidylinositol-4,5-bisphosphate 3-kinase catalytic subunit alpha (PIK3CA), a catalytic subunit of the phosphoinositide 3-kinase (PI3K) complex and an upstream activator of the AKT pathway^61^, which was significantly upregulated in memory B cells (lfsr = 3.18 × 10⁻⁴) and CD4⁺ memory T cells (lfsr = 3.70 × 10⁻²). These changes suggested that phosphatidylinositol (3,4,5)-trisphosphate (PIP₃) generation, a critical step for AKT activation, may be disrupted in this population. In fact, AKT serine/threonine kinase 2 (AKT2), which mediates insulin-stimulated glucose uptake and contributes to anti-inflammatory signaling^62^, showed a positive trend toward upregulation in memory B cells (lfsr = 9.4 × 10⁻²) and was significantly upregulated in NK cells (lfsr = 2.8 × 10⁻²), suggesting altered insulin sensitivity in these cell types. On the other hand, inositol polyphosphate-5-phosphatase A (INPP5A; lfsr = 1.83 × 10⁻⁸) was also one of the most significantly upregulated genes in memory B cells. INPP5A encodes a 5-phosphatase that hydrolyzes PIP₃, thereby acting as a negative regulator on PIP₃^63^implicating a negative feedback mechanism attempting to limit excessive PI3K activity and AKT phosphorylation under chronic insulin exposure.

Furthermore, several transcriptional regulators with dual metabolic and inflammatory roles were dysregulated in memory B cells. Forkhead box O1 (FOXO1; lfsr = 1.6 x 10⁻²) and Sterol regulatory element-binding transcription factor 1 (SREBF1; lfsr = 2.7 x 10⁻²), both downstream of AKT and known to coordinate glucose and lipid metabolism^64^, were upregulated, while Nuclear factor kappa B subunit 1 (NFKB1; lfsr = 1.00 × 10⁻³) and Inhibitor of NF-κB kinase subunit β (IKBKB; lfsr = 2.53 × 10⁻²) showed opposite regulation. Taken together, these findings indicate that insulin-mediated signaling and transcriptional regulation are perturbed in memory B cells of the male HA group.

We also investigated the male HN vs LN contrast with ORA due to the substantial number of differentially accessible peaks observed in that comparison (**Fig. S9**). Interestingly, and similar to the HA vs LN pathway enrichment results, we found consistent over-representation of the term, *‘positive regulation of metabolic process’* for all four cell types tested, including naive B cells (adj. p-value = 1.03 × 10^-4^), CD4+ naive T cells (adj. p-value = 9.31 × 10^-2^), CD14+ monocytes (adj. p-value = 4.6 x 10^-4^), and NK cells (adj. p-value = 5.82 × 10^-7^). In addition, glucose metabolism-related pathways, such as *‘regulation of glucose metabolic processes’* (adj. p-value = 1.08 × 10^-2^) and the *‘insulin signaling pathway’* (adj. p-value = 5.34 x 10^-2^) demonstrated significant enrichment in NK cells. These enriched pathways suggest that chromatin accessibility changes in the male HN group may reflect a history of broad metabolic dysregulation in the absence of asthma. In the case of monocytes, which are well recognized for their capacity to acquire “metabolic memory” following prolonged exposure to stimuli such as hyperglycemia and hyperlipidemia^65,66^, this shared enrichment between HN and HA monocytes suggests a common insulin-driven reprogramming axis, wherein sustained metabolic dysregulation may be consistently epigenetically marked within the myeloid lineage, regardless of asthma status.

To deepen our understanding of the upstream regulators driving the transcriptomic and chromatin-level changes described above, we performed motif enrichment analysis on DA regions from the male HA and HN vs LN comparisons (see Methods: Motif enrichment analysis). At the TF family-level, zinc-finger–containing TFs were the most commonly identified (**Fig. S10A**), likely due to the size of this gene family^67^, however, the family contributing the most significantly enriched TF in both comparisons was the CREB-related (cAMP-response element-binding protein) family (**Fig. S10B**), which includes known mediators of insulin signaling and glucose metabolism^68,69^.

To better target our motif analyses on relevant TFs from the literature, we extracted TFs implicated in inflammatory activation and insulin-responsive regulation from existing resources (5-6 classic TFs for each pathway, see Methods). Inflammation-related TFs, including MYC Proto-Oncogene (MYC), Hypoxia Inducible Factor 1 Alpha Subunit (HIF1A), NFKB1, RELA proto-oncogene, NF-kB subunit (RELA), Interferon Regulatory Factor 7 (IRF7), and Krüppel-Like Factor 6 (KLF6), were found enriched in various cell types across both comparisons (**Fig. 3C**). Notably, these motifs were predominantly detected in B cells for both phenotypes (HA and HN). However, the limited overlap with transcriptional changes in HN B cells suggests that altered motif accessibility does not necessarily translate into active transcriptional responses in this context. Among these TFs, HIF1A motifs were uniquely enriched across all three HA cell types but absent in HN. HIF1A is known for triggering a metabolic shift in cells from oxidative phosphorylation to glycolysis that fuels inflammation^70,71^.

In addition to inflammation-associated TFs, several insulin-responsive TFs displayed widespread motif enrichment. Carbohydrate-Responsive Element-Binding Protein (ChREBP, encoded by the *MLXIPL* gene) binding motifs were found enriched in most cell types across both comparisons, suggesting that insulin-responsive chromatin remodeling is a pervasive feature of high-insulin phenotypes, regardless of asthma status, with the notable exception of HA NK cells. Consistent with pathway-level analyses, CD14⁺ monocytes from both HA and HN males were enriched for insulin-responsive TFs.

### Genotyping and cell-type specific cis-QTL mapping with single-cell multiomic data reveals potential disease pathway

Finally, to attempt to establish an end-to-end mechanistic model of genetic variation influencing our phenotypes, and to fully leverage the TCRS single-cell multiomic dataset, we aimed to perform genotype calling directly from single-cell RNA and ATAC sequencing data for the purpose of QTL mapping. Variant calling and genotyping were performed using Monopogen^72^. In total, we identified 1,980,733 SNPs with high confidence. Comparing the Monopogen-derived genotypes to microarray-based genotype data that was available for 27 of the subjects included in this study, the mean concordance rate for self-comparisons across the two technologies was 83.9% (median: 82.6%), significantly higher (p-value = 2.0 × 10⁻¹⁶) than the mean concordance rate (61.6%; median: 61.4%) for different subjects compared across the two technologies. Principal component analysis (PCA) of the genotypes indicated that the majority of TCRS subjects clustered with European populations or at the interface between European and admixed American ancestries (**Fig. S11C**), consistent with predominant self-reported European ancestry in this population. We also assessed whether single-cell-derived SNPs, particularly those identified from the ATAC modality, are more enriched near regulatory regions. We observed that SNPs derived from single-cell data, especially those from the ATAC modality, were more likely to directly overlap annotated regulatory elements (Fisher’s exact test p-value = 6.66 × 10^-214^, odds ratio = 1.59), indicating better coverage of truly functional regulatory elements (**Fig. S11D**).

Building on this dataset, we next integrated single-cell genotype calls with RNA and ATAC modalities to perform cis-eQTL (expression quantitative trait loci) and cis-caQTL (chromatin accessibility quantitative trait loci) mapping. At a 5% FDR, we identified 2,071 unique genes (“eGenes”; median: 250 per cell type), and 7,063 unique ATAC peaks (“caPeaks”; median: 568 per cell type) with their quantitative measurements significantly associated with proximal SNP variation (**Fig. S12A**). From the opposite perspective, a median of 4,212.5 SNPs per cell type were associated with gene expression in our QTL analysis, while 4,919.5 SNPs per cell type were associated with chromatin accessibility. A median of 276.5 SNPs per cell type were associated with both features simultaneously (**Fig. S12B**).

To prioritize genetic loci potentially contributing to the high-insulin asthmatic (HA) phenotype, we overlapped loci where genetic variation was associated with both gene expression and chromatin accessibility (“peak–SNP–gene triplets”) with the DE genes and DA peaks identified in the previous sections. A two-threshold strategy for identifying concordant differential expression and accessibility was implemented to balance biological sensitivity with statistical rigor: for each gene/peak pair within a triplet, we required one test (differential expression or differential accessibility) to meet a stringent threshold (lfsr < 0.1) and the other was required to pass a relaxed threshold (lfsr < 0.5). This strategy was based on the assumption that peak/gene pairs linked by the same SNP are more likely to exhibit biological correlation, which we validated by comparison against random background peak/gene pairs (**Fig. S12C**). Applying this method, we identified 192 significant peak–SNP–gene triplets corresponding to 10 distinct peak/gene pairs (**Fig. S13**). Consistent with earlier findings highlighting memory B cells as a key affected population in male HA subjects, two memory B cell-specific peak/gene pairs - *HLA-DQB1* with chr6:32,636,768-32,637,577, and *AHI1* with chr6:135,497,163-135,498,726 - were found to be linked by shared cis-QTLs and passed the two-threshold filter. Looking more closely at *HLA-DQB1* (**Fig. 4**), which has previously been implicated in asthma pathogenesis^73–76^, we identified five cis-acting SNPs associated with both *HLA-DQB1* expression and chromatin accessibility at the genomic region chr6:32,636,768–32,637,577. By jointly visualizing chromatin accessibility and gene expression within a 100 kb window centered on *HLA-DQB1* in memory B cells from male HA and LN subjects, we observed elevated *HLA-DQB1* expression in the HA group compared to LN (lfsr = 9.10 x 10^-2^; logFC = 0.85). Correspondingly, chromatin accessibility at chr6:32,636,768–32,637,577 trended towards being more accessible in HA samples (lfsr = 0.43; logFC = 0.29), although the signal did not reach statistical significance. Of the five SNPs identified, four were located within intronic regions of *HLA-DQA1*, while one mapped to an intron of *HLA-DQB1*. Examining the five most significant cis-SNPs as representative variants (**Fig. 4B, Fig. S14**), we observed a consistent dosage-dependent relationship in which having more alternative alleles was associated with reduced *HLA-DQB1* expression and chromatin accessibility. Taken together with previous findings, these trends suggest that the alternative allele may drive decreased HLA-DQB1 expression through decreased chromatin accessibility at this site in memory B cells of HA males and could implicate the alternative allele as a potentially asthma-protective variant.

**Figure 4.**
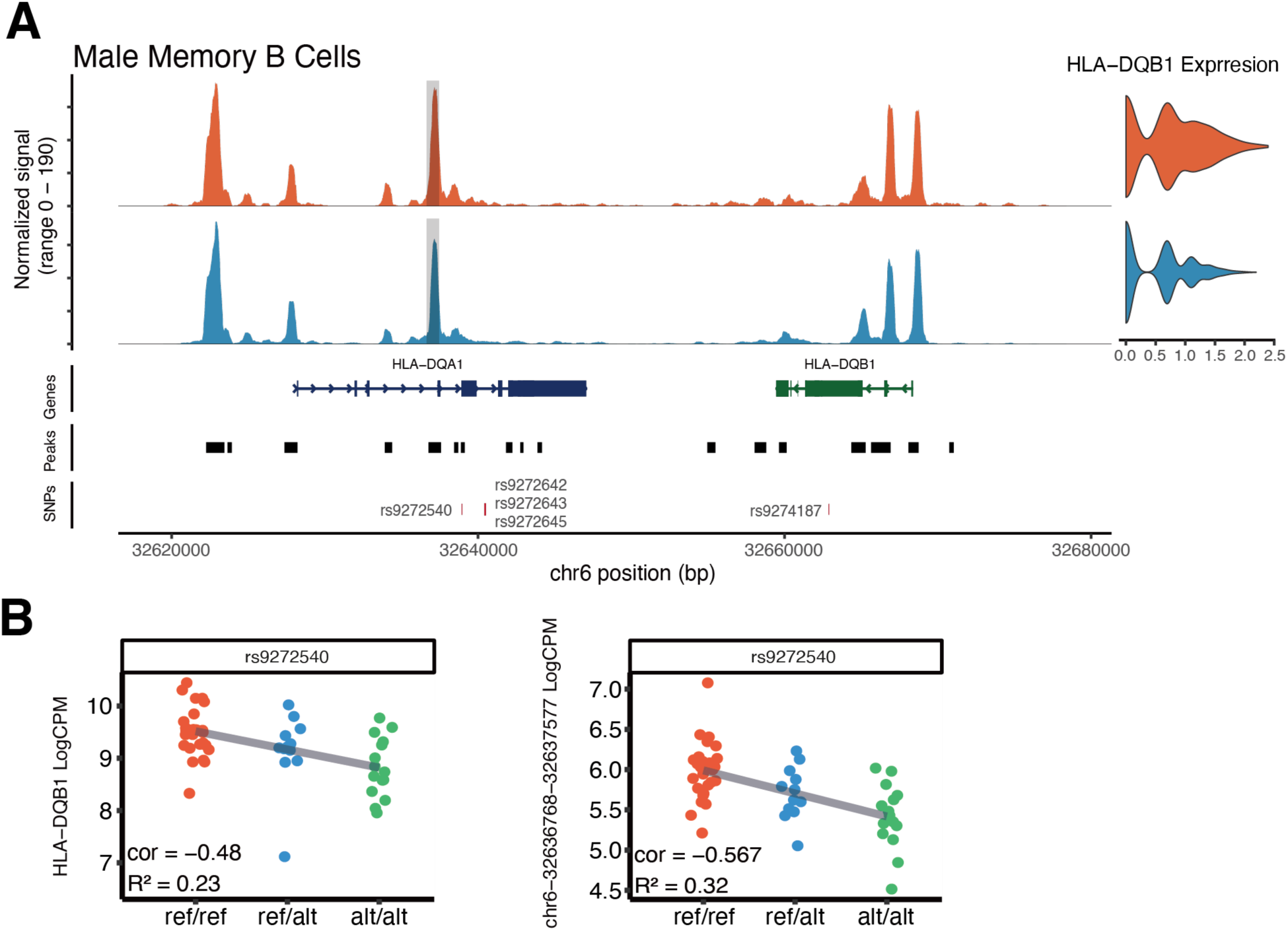
cis-QTL analysis using single-cell multiomic data. **(A)** Integrated visualization of peak accessibility and gene expression across a ±50kb genomic window centered on HLA-DQB1. Peak coverage plot: blue peaks correspond to the LN group, red peaks to the HA group; the gray bar marks the location of the peak at chr6:32,636,768-32,637,577. Gene expression plot: violin plots represent expression levels for LN (blue) and HA (red) groups. Peak location track indicates identified accessible chromatin regions. Gene annotation track displays genes located within the shown genomic range. SNPs track displays loci where five cis-acting SNPs. **(B)** Dot plots illustrating the relationship between genotype and molecular phenotypes for one of the five cis-acting SNPs (rs9272540). The left panels depict normalized expression levels of the HLA-DQB1 gene, while the right panels show chromatin accessibility at the genomic locus chr6:32,636,768–32,637,577. Data points are stratified by genotype and color-coded as follows: reference homozygous (red), heterozygous (blue), and alternate homozygous (green).

Other regions beyond HLA were also implicated by our analysis. The gene-peak pair associated with the largest number of SNPs was “Component of Oligomeric Golgi Complex 5” (*COG5*) and chr7:107,560,885–107,561,285 in NK cells (75 SNPs associated), specifically in the male HA vs LN comparison. While *COG5* itself has not previously been associated with asthma, it plays a role in glycosylation, a process that has been shown to be essential to NK cell function^77^ and has a role in asthma^78^. The gene implicated across the greatest number of comparisons was “Stabilizer Of Axonemal Microtubules 2” (*SAXO2*)*. SAXO2* was affected in naïve CD4⁺ T cells when comparing HA and LN in females, as well as both naïve CD4⁺ T cells and NK cells in males when comparing LA and LN to controls (**Fig. S13**). *SAXO2* is proposed to play a role in microtubule formation and cytoskeletal organization^79^. Like *COG5*, *SAXO2* has not been previously implicated in asthma, however, NK cell function^80^ and regulation of immune cell extravasation^81^ are critical pathways in asthma.

## Discussion

In this study, we developed and validated a cost-effective, scalable, and user-friendly single-cell multiomic profiling method, called mTEA-seq, to investigate how early-life metabolic dysregulation shapes immune cell states later in life. Leveraging a well-characterized birth cohort with longitudinal metabolic and asthma phenotyping, we profiled PBMCs from individuals stratified by early-life serum insulin levels and asthma diagnosis up to age 40. Our observations highlight three key points that together support an insulin-associated asthma endotype.

First, the molecular phenotypes we observe are remarkably sex-specific. The vast majority of identified differences between “case” group and control samples are from comparisons of the male subjects. We formally tested this and were able to identify a significant sex interaction term for a large fraction of these observations, in spite of the conservative nature of such a test. We believe that these findings are highly consistent with the known biology of both asthma and insulin signaling, which are both documented to have pervasive sex-specific effects^48,49^. This sex bias was consistent across all three “case” groups: those with asthma, but no documented insulin dysregulation; those with early-life elevated insulin, but no asthma; and those with both early-life elevated insulin and an asthma diagnosis. To the degree that our results support an early-life insulin dysregulation-driven endotype of asthma, they suggest that this is likely to primarily affect males.

Second, we observed an ‘uncoupling’ of epigenetic and transcriptomic rewiring in comparing the male subjects with early-life elevated insulin and no asthma (HN) to those with early-life elevated insulin and an asthma diagnosis (HA). The HN group reflected epigenetic changes across many cell types, but relatively few transcriptional changes, suggesting the possibility of a primed state that has not been activated. On the other hand, the HA group exhibited both transcriptional and epigenetic changes. The epigenetic changes in both groups were associated with metabolic pathways, while the transcriptional changes in the HA group were largely associated with inflammatory pathways. The observation of epigenetic changes in response to past metabolic states is consistent with the literature on metabolic memory^65,82,83^ and trained immunity^84,85^. Remarkably, the degree to which epigenetic changes were shared between these two groups was extremely cell-type-dependent. CD14+ monocytes show a large degree of sharing in their epigenetic rewiring in the HA and HN states, while NK cells show rewiring that is almost totally independent between HA and HN, and cell types like memory B cells are somewhere in between. While the consistent effect of high insulin on monocytes may help to explain why monocytes are so often studied in the context of metabolic memory and trained immunity^65^, we speculate that the discrepancy in NK cell responses is most likely to help explain the difference between individuals with early life high insulin who went on to develop asthma and those who did not. It is established in the literature that NK cells can form long-lived trained memory populations that can have distinct effects upon subsequent stimulation, depending on the context of the training^86^. Key regulators that are differentially enriched among affected targets in HA NK cells include KLF6 and HIF1A, both of which have established roles in insulin signalling^87–91^, are pro-inflammatory^92–95^, regulate the transcription of each other^96,97^, and have been implicated in NK cell trained immunity^92,98^, suggesting that they may play vital roles in translating aberrant insulin signalling into an overactive inflammatory state in NK cells in the high-insulin asthma endotype. On the other side of the spectrum, HN NK cells showed enrichment of IRF7 and ChREBP, encoded by the *MLXIPL* gene, suggesting a retained anti-viral capacity^99^ and metabolic homeostasis^100^.

Finally, our study highlights a particularly long time between the measurement of the risk factor (insulin levels at age 6) and the detected molecular consequences (transcriptional and epigenomic differences in PBMCs collected at age 40). These results suggest that early-life metabolic risk exposures could leave durable, measurable imprints on the adult immune system. Previous studies have documented effects of trained immunity that have lasted several months in humans^101–103^ and over a year in mice^104^. Given the discrepancy in lifespans, these are strikingly different time scales, implying that the time frame for trained immunity persistence might be considerably longer than has yet been documented in humans. While our data are not necessarily conclusive evidence of metabolic memory, they propose a much longer benchmark for proposed maintenance of memory in immune cells and are consistent with a recent study showing that elevated insulin levels can establish durable insulin resistance in several cell types and a mouse model^105^. It is also possible that the early-life nature of the insulin exposure in our study population represents the disruption of a critical window of development. An alternative interpretation that we cannot exclude, however, is that the early-life environmental factors that lead to elevated insulin levels are remarkably persistent over the life course of our subjects, and therefore the elevated insulin at age 6 simply serves as an early biomarker of lifelong struggles with maintaining normal insulin levels. Future research will have to disentangle these possibilities.

Nevertheless, several limitations of this study are worthy of discussion. These include the fact that despite the use of multiplexing to enhance throughput, the study remains constrained by a relatively modest sample size compared to the inherent interindividual variability of human samples. This may have limited our statistical power in detecting differential features in disease phenotype groups, especially given the observed sexual dimorphism. With regard to the strong sex-specific effects we observed, the underlying biological reasons for male-biased molecular alterations remain unclear and need further investigation in larger, independent cohorts. Further, although we implemented sophisticated statistical modeling to control for repeated measures and technical covariates, residual batch effects or unmeasured confounders cannot be entirely excluded. In addition, while we achieved a mean concordance rate of 83.9% between single-cell-derived genotypes and microarray-based genotyping, which is sufficient to reliably identify the correct subject, the sparse and biased coverage characteristic of single-cell sequencing still constrained genotyping accuracy at specific genomic loci (**Fig. S15**). In particular, regions of low or uneven coverage may yield less reliable SNP calls, especially in the absence of high-confidence linkage disequilibrium-based imputation. These limitations may have reduced the resolution of cis-QTL mapping and impaired the interpretability of SNP-specific analyses in a subset of individuals. Lastly, while we have identified candidate regulatory variants and pathways, future functional validation (e.g., CRISPR perturbation or allele-specific assays) will be necessary to confirm inferred causal relationships. In spite of these caveats, we believe our data are consistent with and expand upon existing knowledge in the realm of metabolism and immune function.

In summary, this study established a cost-effective framework for population-scale single-cell multiomics and revealed immune and metabolic alterations linked to early-life insulin dysregulation. The gene expression and chromatin accessibility differences we identified underscore the lasting impact of early-life metabolic states on adult immune function. The convergence of insulin-driven metabolic reprogramming and inflammatory activation in males supports an insulin-associated asthma endotype that links metabolic history to immune dysfunction. These findings highlight how sex-specific metabolic–immune interactions may shape this asthma endotype and provide a framework for investigating how early metabolic environments influence asthma development later in life.

## Data Availability

Raw sequencing data, genotype vcf files, and processed feature-count matrices will be provided through dbGaP.

## Code Availability

Code will be made available through GitHub.

## Acknowledgements

We thank the families and participants of the Tucson Children’s Respiratory Study for their longstanding commitment, and the research nurses, especially Silvia Lopez, for their dedication. We thank Dr. Marilyn Halonen for thorough and insightful comments on the manuscript. We are grateful to Dr. Hao Zhang, Dr. Holly Welfley, Dr. Sydney Harding, and members of the Romanoski Lab at the University of Arizona for their valuable advice and support. We thank Jonathan R. Galina-Mehlman, of the University of Arizona Genetics Core, Dr. Daniel Laubitz, of the PANDA Core for Genomics and Microbiome Research, Mark Curry, of the University of Arizona Cytometry Core Facility, and Sara Willis, of University of Arizona Information Technology Services, for their expert technical assistance. We also acknowledge the University of Arizona High Performance Computing core for providing computational resources.

## Methods

### Insulin level classification and asthma diagnosis

As previously reported^36^, non-fasting insulin was measured by multiplex (Human Multi-Analyte Profile panel version 1.6, Myriad Rules Based Medicine, Austin TX) on all individuals of the cohort for whom serum collected at the age 6 visit was available for analysis (n=383). Insulin analyses were limited to measurements in a single batch of serum samples measured over 7 plates (n=342). Results for insulin from RBM assays were corroborated by ELISA (R&D Systems, Minneapolis, USA) in a randomly selected subset of the same samples used for the RBM assay. Insulin levels at age 6 were divided into quartiles, separately by sex. High insulin was defined as the top quartile; low as the bottom three quartiles. We used these cohort-wide definitions to define participants in this sample.

Physician-diagnosed active asthma at age 6 and in subsequent surveys through age 40 was defined as a physician diagnosis plus active symptoms in the past year. Participants were classified as ever asthma if there was active asthma at any survey, and never asthma if no completed survey documented asthma diagnosis.

### Blood sample processing

Peripheral blood was collected by venipuncture and processed within 24 hours. PBMCs were separated over lymphocyte separation media using standard techniques. Cells were immediately frozen in RPMI + 40% FBS + 10% DMSO and cryopreserved until the time of assay.PBMC samples and mTEA-seq protocol The 4 PBMC test samples for mTEA-seq method development were obtained from STEMCELL Technologies and cryopreserved immediately upon arrival. A detailed protocol for sample processing and library preparation has been documented and made publicly available on protocols.io (https://www.protocols.io/edit/mtea-seq-edited-dj2u4qew). Test samples were processed as described below for TCRS samples.

### TCRS PBMC sample preparation

Samples were processed in two batches that included repeated profiling of most of the asthmatics and high insulin subjects, which allowed us to assess robustness of the measurements. To achieve balanced representation across the four phenotypic groups, the initial batch was intended to include 10 individuals from each group. The second batch was intended to include 20 new control individuals along with the case group samples profiled in the first batch. Batch 1 included 39 samples, while Batch 2 included 50. In Batch 1, samples were randomly assigned to five groups, ensuring balanced representation of four predefined phenotypic categories (LN, HN, LA, HA). Groups 1–4 each contained eight samples, and Group 5 included seven samples. Batch 2 followed a similar randomization strategy, allocating 10 samples per group with balanced phenotypic distribution. For single-cell library generation, Batch 1 groups were processed using two 10x Genomics channels per group, while Batch 2 utilized three channels per group to enhance data throughput. A summary of sample identifiers, phenotype assignments, and batch/channel mapping is provided in Table S1.

#### PBMC staining and sorting

Frozen PBMCs isolated from subjects were warmed up at 37 °C for 2 minutes and diluted with 9-volumes of prewarmed AIM-V (Gibco #12055091), then subjected to centrifugation. The cell pellets were washed with 10 ml of ice-cold Cell Buffer (0.2% BSA in PBS buffer), pelleted by centrifugation, and followed by resuspension with 1 ml of cell buffer. All pelleting of samples was conducted at a speed of 400 x g for 5 minutes at 4 °C, in a swinging bucket centrifuge. Cell numbers were determined by trypan blue staining and counting using a Countess Automatic Cell Counter (Invitrogen).

One million PBMCs from each subject were stained with 0.001 X FVS-510 (BD #564406), depending on the volume from the previous step, in 100 ul of PBS for 15 minutes at room temperature and then pelleted to change the buffer to 100 ul Staining Buffer (2% BSA in PBS), followed by 10% TruStain FcX (Biolegend #422302) staining at 10 minutes at room temperature. After the blocking reaction, 300 ul of Staining Buffer, 3.8 ul of hashtag antibody (TotalSeq™-A anti-human Hashtag antibody, Biolegend), and each 5 ul of antibodies against CD45 (FITC anti-HumanBiolegend, #304038), CD14 (APC anti-human, Biolegend #301808), CD15 (PE Mouse anti Human, W6D3, BD #562371), and CD16 (PE/Cyanine7 anti-human, Biolegend #302016) were added to each sample and incubated for 30 minutes on ice. Unbound antibodies were washed out with 4 ml of ice-cold AIM V twice, and the stained cell pellet was resuspended with the Staining Buffer for sorting by BD Aria II.

Debris was removed by gating based on side and forward scatter area (SSC-A and FSC-A), and single cells were isolated by FSC-H vs FSC-A to exclude doublets. Dead cells were identified as FVS-510-positive and excluded from downstream analysis. White blood cells were enriched by positive selection for CD45+ populations. Neutrophils were removed using a high SSC-A signal and high CD15 expression, as the extracellular matrix and high RNase content can negatively impact single-cell data quality. To prevent unintended exclusion of monocytes, which also express CD15, the expression patterns of CD14 and CD16 were assessed to confirm monocyte retention. For each experimental group, a minimum of 3 million cells was targeted for sorting to ensure sufficient input material for permeabilization and library preparation. Sample contributions were balanced across groups to the best of our ability to reduce bias in downstream analyses.

#### Library Construction

One million sorted and pooled cells were permeabilized with 100 ul of Permeabilization Buffer containing 0.01% Digitonin in the Washing Buffer (20 mM Tris-HCl (pH7.5), 150 mM NaCl and 3 mM MgCl2, and 1U/ul Protector RNase inhibitor) on ice for 5 minutes. Permeabilization was terminated by adding 1ml of Washing Buffer and followed by centrifugation. The cell pellet was resuspended with 25 ul of Washing Buffer and the cell number was determined by trypan blue staining.

A total of 40K cells was subjected to 10X Chromium Next GEM Single Cell Multiome ATAC + Gene Expression profiling according to the TEA-seq paper (original TEA-seq protocol on protocols.io) with separate amplification of cDNA and HTO samples. The HTO indexing PCR was done directly using the pre-amplification product by Kapa HiFi DNA polymerase master mix (Roche #07958927001) and cleaned up with 1.6X SPRI beads. The amplified cDNA without spike-in additive HTO Primer was cleaned up with 0.6X SPRI beads. All PCRs were monitored on real-time PCR by SYBR Green I and the size distributions of final libraries were confirmed by TapeStation analysis. All detailed methods are described at: https://www.protocols.io/edit/mtea-seq-edited-dj2u4qew.

### TCRS single-cell multiomic data cleanup

#### Library sequencing and Fastq generation

TCRS libraries (cDNA, ATAC, and HTO) were sequenced on an Illumina NovaSeq 6000 system at the PANDA Core Genetics and Microbiome Research Facility at the University of Arizona, using the following read configuration: R1=50 bp, Index1=24 bp, Index2=10 bp, and R2=90 bp. cDNA and HTO libraries were demultiplexed using bcl2fastq. Demultiplexed ATAC and cDNA FASTQ files were processed using 10x Genomics CellRanger arc v2.0.1^5^ ‘cellranger-arc count’ to generate co-assay count matrices and genome alignment BAM files. To accommodate our high-throughput workflow, we bypassed the default cell limit of the 10x Genomics CellRanger arc v2.0.1 program by modifying an internal cell number cap, MAXIMUM_CELLS_PER_SPECIES, from 20,000 to 50,000. Raw count matrices generated by 10x Genomics CellRanger arc v2.0.1 were imported into RStudio for quality control using Seurat^106–110^.

#### Cell identification and quality filtering

Cell barcodes were classified as valid cells when their UMI counts and ATAC fragment density were different from background (empty droplets) density. The thresholds were determined via k-means clustering (k=2), with the cutoff defined as the background cluster centroid plus 5% of the distance between the two centroids (**Fig. S16**). Valid cell barcodes passing both cDNA and ATAC thresholds were retained as “whitelist” barcodes. These barcodes were used with CITE-Seq-Count to generate HTO UMI matrices, by setting no correction for umi (--no_umi_correction) and max 2-edit distance for HTO tag (--max-error 2).

For each experimental channel, RNA Seurat objects were created, and HTO counts were added as a separate ‘HTO’ assay. HTO identities were assigned using Seurat’s HTODemux function (positive.quantile = 0.99). Cells were filtered based on the following criteria: (1) number of detected genes should be within 200–2,500 range; (2) mitochondrial read percentage <20%; and (3) positive and unambiguous HTO detection. For the purpose of creating initial Seurat object for quality control and comparing mTEA-seq ATAC data quality with TEA-seq, we extracted an orthogonal single-cell PBMC peak set from a single-cell multiomic data set, ‘pbmcMultiome.SeuratData’, available from ‘SeuratDisk’. We generated ATAC feature count matrices using this orthogonal peak set, only considering cells passing cDNA and HTO quality controls described above. We then calculated FFiP and TSS enrichment scores using Signac^111^ functions ‘FRiP’ and ‘TSSEnrichment’. cDNA and HTO controlled cells were further controlled on ATAC matrices with FFiP > 25% and TSS enrichment > 2.

Following HTO demultiplexing, the median heterotypic (droplets with more than one HTO observed) multiplet rates were 28.2% for Batch 1 and 23.5% for Batch 2. An additional 3.53% (1/8 of observed collision rate) and 2.35% (1/10 of observed collision rate) of homotypic doublets (same-hashing multiplets) for batch 1 and batch 2 were estimated to be undetectable by HTO-based methods. Despite their low prevalence, we addressed this issue using Scrublet setting an expected doublet rate of 3% to identify and remove cells with aberrant transcriptomic profiles indicative of multiplet artifacts.

In addition to single-cell quality control, we also excluded ten samples(6 unique subjects) from downstream analysis due to cell viability <50% detected during sorting (**Fig. S1B**). One sample was excluded at the stage of FACS sorting due to severe clumping. In addition, one sample from Batch 1 was inadvertently removed from the analysis pipeline because it was mis-labeledas a low viability sample. However, this sample was also included in Batch 2, and so we elected not to re-integrate the data excluded in error.

After stringent quality filters, including cDNA, HTO, ATAC, Scrublet, and FACS viability, a total of 279,888 high-confidence and high-quality single cells of 55 unique subjects were retained. However, one additional sample was later identified as a potential label-swap sample (based on discordance between microarray and single-cell genotype calls). Therefore, this sample was removed from all analyses and 272,003 PBMCs remained, accounting for 54 unique subjects, for the differential analysis.

### TCRS Data integration and label transfer

Robust single-cell data integration is critical for accurate cell type annotation. We implemented the following multi-modal integration strategy: 1) Independent modality integration: cDNA and ATAC data were integrated separately to address modality-specific technical biases. 2) Cross-modality synthesis: Integrated ATAC and dimension reduction information were passed to the harmonized cDNA Seurat object, enabling the creation of a unified coassay object.

#### cDNA integration

We first merged batch 1 and batch 2 cDNA plus HTO Seurat object lists. We then used the RunHarmony function from Harmony package reducing the effects of batch, different 10X channels, sex, and disease state. The Integrated cDNA object was run through the regular Seurat pipeline, including normalization, scaling, PCA reduction, identifying clusters, and UMAP projection. Dimension reduction UMAP was inspected to ensure cell clustering was not dominated by any technical effects.

#### ATAC integration

In order to retain disease state related epigenetic modifications across cell types, we employed an ‘iterative peak calling and counting’ integration strategy. In the first iteration, ATAC count matrices from batch 1 and batch 2, both generated on orthogonal peaks, were merged, and new peak sets were independently called for each of the phenotypic subgroups (LN, HN, LA, HA) using MACS2. Raw peaks were filtered to exclude regions overlapping the ENCODE hg38 genome blacklist and overlapping regions were merged. Filtered phenotypic peaks were used to regenerate new fragment count matrices, and ATAC Seurat objects were integrated across batches and 10X channels using Harmony integration reducing the effects of 10X experimental channel, batch, disease state, and sex. Integrated ATAC embeddings and assays were combined with the harmonized cDNA object (corrected for channel, batch, sex, and disease state via Harmony) to construct a multi-modal co-assay object. Multimodal neighbors were detected using FindMultiModalNeighbors, incorporating cDNA Harmony reductions (dims 1–30) and ATAC Harmony reductions (dims 2–30). We used an annotated PBMC dataset^107^ to perform label transfer primarily based on transcriptional profile as demonstrated by Signac vignettes - joint RNA and ATAC analysis: 10X multiomics. This iteration prioritized retention of phenotype-associated chromatin features.

In the second iteration, peaks were re-called by first aggregating cells by cell type annotations from the first round (not by individual or disease class) to refine accessibility features linked to cell states. We only utilized confidently labeled cells for calling peaks, with prediction score > 0.5 (235,680 cells of total 279,888 cells, or 84.21% overall). New peaks underwent identical filtering mentioned above (i.e. blacklist exclusion, interval merging) and were used to regenerate ATAC count matrices. These matrices were reintegrated using the same workflow (Harmony integration) and merged with the harmonized cDNA object to generate a final multi-modal co-assay. This iterative approach enhances resolution of epigenetic heterogeneity across cell types while preserving phenotype-specific chromatin signatures, enabling robust integration of transcriptional and chromatin accessibility landscapes.

#### Label transfer and annotation Cleanup

Single-cell label transfer is a fast and efficient method for annotating single-cell datasets by leveraging pre-labeled references, allowing researchers to bypass the challenges of selecting cell markers and fine-tuning clustering granularity, especially for closely related cell types. However, because label transfer assigns cell identities based on the nearest match in the reference dataset, mislabeling occurs when technical noise between the query and reference datasets is substantial. In response to those potential pitfalls, we selected a publicly available single-cell multiomic data set^107^ that utilizes the same 10X platform as our TCRS data, thereby minimizing platform-related discrepancies. We then refined the annotations by collapsing transferred labels based on the majority vote within clusters defined by the TCRS data, setting the WNN resolution to 2, using the first 20 principal components (PCs) for cDNA, and 2:20 PCs for ATAC. Majority rule was adapted to handle edge cases in two scenarios: 1) when label collapsing resulted in the loss of rare cell types (e.g., Treg, dnT, or gdT), we subclustered the target cluster by increasing the dimensionality reduction to 30 components, then applied majority voting to recover the rare cell types; 2) when the dominant cell type was ambiguous (i.e., < 20% difference in proportion between the first and second most frequent cell types), we similarly subclustered the target cluster by increasing the dimensionality reduction to 30 components, and then used majority vote to resolve the ambiguity. Finally, we validated our cell type annotations by checking expression of cell type markers and visually comparing them against the original reference datasets across a spectrum of cell type markers (**Fig. S5**).

### Cell type proportion differential analysis

To evaluate differences in immune cell type proportions across phenotypic groups, we conducted a differential cell proportion analysis stratified by sex. We quantified the number of cells assigned to each predicted cell type and computed the relative proportion of each cell type within its respective sample batch. In cases where a sample had repeated measurements, cell type proportions were averaged across replicates. Comparisons were performed between case groups and control groups (i.e., LA vs LN, HN vs LN, and HA vs LN) using the Student’s t-test. Only groups with at least three independent subject measurements were included in each comparison. To correct for multiple testing across cell types within each sex, p-values were adjusted using the Benjamini–Hochberg (BH) procedure. Results were filtered to retain comparisons with adjusted p value < 0.05.

### DE gene and DA peak differential analysis Using mashr with limma Covariance Matrices

To identify phenotype-associated differential genes and peaks within each sex, we conducted an analysis integrating pseudobulk aggregation, linear modeling, and multivariate adaptive shrinkage powered by mashr. mashr is a Bayesian framework that estimates and shrinks effect sizes toward data-driven covariance structures shared across conditions. The method requires as input gene-(or peak-) specific estimated effects and their standard errors from each condition, along with the empirical correlations among those effects. Specifically, in our analysis, each “condition” corresponds to a phenotype–sex contrast (e.g., HA vs LN in males or females), and mashr models the joint distribution of these effects to identify shared or condition-specific differential patterns while improving power and controlling false discovery.

Single-cell RNA-seq data were preprocessed by removing one sample, identified as a potential label-swapping outlier during genotyping validation. Metadata covariates, including sample ID, batch (b1/b2), insulin level (L/H), asthma status (N/A), sex, and derived interaction terms (group: phenotype-sex combination), were factorized to ensure consistent modeling. Cell types with median counts < 30 per sample or fewer than three unique subjects per phenotype-sex group were excluded. These thresholds mitigate overdispersion in low-count data and ensure stable variance estimation, while also guaranteeing minimum 3 biological replicates per phenotype-sex group. Pseudobulk counts were generated by aggregating cells per subject-batch-cell type combination, followed by library size normalization, with the subject-specific library size calculated using the TMM method from edgeR^112,113^ and mitochondrial gene removal^112,113^.

Using limma-voom^114^, we modeled gene expression with a design matrix incorporating phenotype-sex combinations and the batch label of samples to control for batch effects. Contrasts were defined to compare high-insulin (H), asthma (A), and their combinations against low-insulin non-asthma (LN) baselines, stratified by sex. As some samples were repeatedly measured in batch 1 and batch 2, ignoring within-subject correlation can inflate false positive rate. To account for within-subject correlation, we set the ‘block’ argument in voomLmFit as subject ID toestimate correlations between observations from the same subject, which were in turn used to adjust standard errors of phenotype effect estimates. The actual usage is as: ‘fit <- voomLmFit(counts = dpb, design = design, block = sample_id, sample.weights = TRUE)’

When using mashr, it is crucial to account for the correlation structure among contrasts to ensure valid statistical inference. Shared sources of variability (e.g technical noise) can induce dependencies in the observed effect estimates, even if the true effects are independent. Ignoring this correlation can lead to inflated false discoveries or underestimation of effect sharing. To address this, we derived covariance and correlation matrices for each gene from the limma-voom model, which naturally captures the dependency between contrasts through its linear modeling framework. For a given gene *g*, the weight matrix *V*_*g*_ was calculated as:

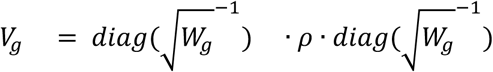

where *V*_*g*_ represents voom precision weights, and *ρ* is the intra-block correlation matrix estimated from repeated measures (via ‘block = sample_id’). The contrast-specific covariance was then computed as:

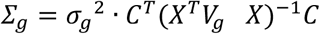

where *X* is the design matrix, *C* is
the contrast matrix, and
*σ*_*g*_^2^is the residual variance. The covariance matrix was then converted into a correlation matrix using R ‘cov2cor’ function. To resolve discrepancies due to floating-point precision in correlation matrix, minor asymmetries in correlation matrices (i.e., |*Cor*_*g*[*i*,*j*)]_ − *Cor*_*g*[*j,i*]_| < 10^-6^) were corrected by symmetrizing:

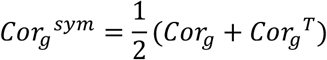

Effect sizes were extracted from the resulting table generated by the limma function ‘contrasts.fit’ and ‘eBayes’. Per gene effect sizes and standard errors, along with the correlation matrix calculated by limma were used as inputs for mashr. An effect was determined as significant if the local false sign rate returned by mashr is < 0.05, which controls the probability that the sign of an effect size is incorrectly estimated below 0.05^115^. The same pipeline was applied to RNA and ATAC pseudobulk count matrices.

#### Evaluation of sex-specific transcription response using interaction modeling

To evaluate whether transcriptional changes associated with the high-insulin asthmatic (HA) phenotype differed between sexes, we performed a sex–phenotype interaction analysis using pseudobulk expression data. We performed the exact same data aggregation, filtering, normalization, and repeated measurements blocking strategy as mentioned above, however the linear model incorporated additional phenotype:sex interaction term. The interaction term tested whether the log(fold-change) between case and control differed significantly between males and females. Genes showing nominally significant interaction effects (p-val < 0.05) were classified as exhibiting sex-specific regulation.

#### Pathway over-representation analysis

Pathway over-representation analysis was performed separately for ATAC-seq peaks and RNA-seq derived gene lists using the HOMER^116^ (Hypergeometric Optimization of Motif EnRichment). Significant differentially accessible ATAC peaks were annotated to nearby genes based on proximity to transcription start sites (TSS) using the human reference annotation (*TxDb.Hsapiens.UCSC.hg38.knownGene)* and the R package ChIPSeeker, allowing a maximum distance of 100 kb from the TSS. Differentially expressed gene lists were converted to Entrez IDs using the clusterProfiler package and the human annotation database (*org.Hs.eg.db*). Comparisons with fewer than 100 Entrez IDs were excluded from enrichment analysis due to low statistical power. The resulting Entrez ID list was then provided as input to HOMER’s findGO.pl to identify enriched pathways. The enrichment score was calculated as the ratio of the proportion of target genes annotated to a given term to the proportion of all background genes annotated to the same term. Finally, p-values from the pathway over-representation analysis were corrected for multiple testing using the BH method. Only significant (adjusted p-value < 0.1) GO-Biological Process and KEGG terms were used for the visual representation.

#### Motif enrichment analysis

To identify transcription factors potentially mediating chromatin accessibility changes, we defined an active motif as one whose binding region was enriched among differentially accessible peaks and whose corresponding transcription factor gene showed expression differences between conditions. DA peaks (lfsr < 0.05) were extracted separately for males in the HN vs LN and HA vs LN contrasts. For each cell type with at least 100 DA peaks, we used FindMotifs from Signac to identify enriched transcription factor binding motifs within the ATAC modality, using background peaks matched for GC content and sequence length. Motifs with adjusted *p*-value < 0.001 and fold enrichment > 1.5 were retained. To evaluate transcriptional activity of these transcription factors, we used FindMarkers from Seurat on the corresponding RNA modality, limiting results to genes expressed in at least 1% of cells and showing greater than 1.5-fold expression difference between case and control groups. Transcription factors meeting both criteria were considered potential regulators. Motif families were annotated with the MotifDb package. Transcription factors were extracted from the MSigDB *HALLMARK_INFLAMMATORY_RESPONSE* gene set and prior studies^64^.

### Genotyping with single-cell multiomic data

For the purpose of discovering genome loci actively regulating identified chromatin accessibility and gene expression differences between cases and controls, we utilized our single-cell multiomic data to call genotypes on the TCRS cohorts.

#### Processing BAM files

cDNA and ATAC BAM files (one per 10X Genomics channel) were first quality controlled using the ‘-F 3844’ and ‘-q 10’ filters in samtools^117^ to remove secondary alignment reads, optical PCR duplicates, any reads that may have failed Illumina quality checks, and reads with a mapping quality < 10 by. We then extracted 10X cell barcodes associated with TCRS subjects and split QC filtered BAM files into individual subject BAM files by using the cell barcodes and prior HTO-based assignments. Deduplication of reads in individual cells was also done simultaneously by filtering on 10X barcode (CB Tag) and UMI (UB Tag) for cDNA or 10X barcode (CB Tag) and genome positions for ATAC BAM files. The resulting BAM files for each subject across 10X channels were further merged into one BAM file for each individual per batch. Because some samples were repeatedly measured in both batches, to avoid significant sequencing depth bias for genotyping, instead of merging batches, we selected the BAM file that had more reads. On average, each subject achieved a mean coverage of 1.18X (median: 0.76X) across the 22 autosomes. Differences in coverage between RNA and ATAC modalities were also quantified across chromosomes (**Fig. S15**).

#### Processing 1000 Genome Project reference

We downloaded a high-coverage 1000 Genomes Project^118^ (1KGP) phased WGS panel consisting of high-quality SNV and INDEL calls across 3,202 1KGP samples from 1KGP FTP server (https://ftp.1000genomes.ebi.ac.uk/vol1/ftp/data_collections/1000G_2504_high_coverage/working/20201028_3202_phased/). The GRCh38 genome reference was obtained from the 1KGP FTP server (https://ftp.1000genomes.ebi.ac.uk/vol1/ftp/technical/reference/GRCh38_reference_genome/).

For each chromosome, the 1KGP VCF records were checked against the GRCh38 genome reference and multi-allelic sites were broken into multiple records with bcftools^119^. Subsequently, variant IDs were replaced with a composite identifier based on chromosome, position, and reference and alternate alleles. A second normalization step was then applied to remove duplicate records and convert the output to BCF format. Converted BCF files were next transformed into PLINK binary format using plink^120^. To reduce redundancy due to linkage disequilibrium and to focus on common variants, we performed minor allele frequency filtering (MAF > 0.1) and linkage disequilibrium (LD) pruning using a sliding window approach (window size = 50, step = 5, variance inflation factor threshold = 1.5). Finally, the list of pruned plink binary files was compiled, and the datasets were merged across chromosomes using plink’s merge-list.

#### Monopogen germline variants detection and imputation

Single-cell data sparsity and uneven coverage pose significant challenges for accurate variant calling and subsequent imputation. To address these issues, we employed an established tool, Monopogen^72^, a tool specifically optimized for single-cell sequencing, which harnesses linkage disequilibrium information from external reference panels to improve variant detection accuracy compared to conventional bulk-based methods such as FreeBayes^121^, GATK^122^, samtools^117^, and Strelka2^123^. Using the “monopogen preProcess” and “monopogen germline” functions, BAM files for each subject were filtered and germline variants were called on the 22 autosomes. Following variant calling, the resulting VCF files were subjected to quality filtering with bcftools, retaining only those variants with a genotype score (squared correlation of dosage or DR2) of at least 0.8. These high-quality variants were then imputed using Beagle.22Jul22.46e^124^, which integrates chromosome-specific genetic maps and reference VCF datasets to impute missing genotypes. After imputation, the VCF files were filtered with DR2 > 0.8. Finally, the imputed genotypes were cross-referenced with pruned 1KGP (see Methods: Processing 1000 Genome Project Reference) to keep only MAF>0.1 and valid SNPs.

#### Microarray genotype imputation

Microarray genotype data were first processed and converted from the GRCh37 to the GRCh38 reference coordinates. The GRCh37 reference VCF file was filtered to remove unconventional chromosomes and INDELs before being lifted over to GRCh38 using GATK’s LiftoverVcf tool. The lifted-over VCF files were then normalized against the GRCh38 reference genome and only biallelic variants were retained. To address missing genotypes, we used Beagle to impute SNPs, leveraging haplotype information from the 1KGP reference panel. Variants with a DR2 < 0.8 were excluded. The remaining SNPs were intersected with a normalized and pruned 1KGP PLINK reference panel (see Methods: Processing 1000 Genome Project Reference) and used for super-population principal component analysis with ‘plink --pca’.

#### Comparing single-cell genotype results with microarray genotype

To assess the accuracy of genotype calls from single-cell sequencing data, we compared single-cell-derived genotypes to array-based genotypes obtained for an overlapping set of subjects. SNPs with MAF < 0.1 in either genotyping set were removed. We then identified intersecting SNPs between the Monopogen-derived genotype dataset and the microarray genotype dataset. SNP IDs were harmonized by reassigning variant identifiers to the format CHROM:POS:REF:ALT. SNPs common to both platforms were retained for subsequent analysis. For each subject’s microarray-based genotypes and each subject’s single-cell-based genotypes, the fraction of matching genotype calls across all intersecting SNPs was calculated. The concordance rate was defined as the number of matching genotypes divided by the total number of SNPs compared, expressed as a percentage. Notably, this analysis identified one subject (**Fig. S11A,B**) with an anomalously low concordance rate between single-cell and microarray genotypes (60.1%), suggesting possible sample swapping or mislabeling, who was therefore excluded from the earlier differential analyses (resulting in 54 subjects), though we retained them for the purpose of QTL mapping.

To evaluate whether single-cell derived SNPs are more proximal to regulatory elements, we calculated the genomic distance between SNPs and reference validated regulatory elements using bedtools. Variant calls were obtained from four sources: single-cell multiomic data (combined ATAC and RNA), single-cell RNA modality, single-cell ATAC modality, and imputed microarray-based genotyping. The reference regulatory elements were obtained from the UCSC Genome Browser (hg38 assembly) by downloading the *refSeqFuncElems* track from the "Regulation" group (https://genome.ucsc.edu/cgi-bin/hgTables?db=hg38&hgta_group=regulation&hgta_track=refSeqFuncElems). For each dataset, SNPs were classified as "near" if they fell within ±1,000 base pairs of a regulatory element. We compared the proportion of "near" and "not near" SNPs between single-cell genotyping methods and microarray data using Fisher’s exact tests. one-sided p-values and odds ratios with 95% confidence intervals were reported.

#### cis-QTL analysis & joint analysis of cis-QTL loci with differential genes or peaks

cis-eQTL and cis-caQTL analyses were performed separately for each cell type using the MatrixeQTL^125^ R package in linear model mode. For RNA and ATAC expression data, only samples selected for monopogen-based genotyping were included; therefore there were not any subjects with repeated measures in this analysis. Normalized, log-transformed counts per million (logCPM) expression matrices (for RNA and ATAC, respectively) were tested against genotype dosage data using a cis-window of 100 kb around each gene or peak. Batch effects were included as covariates in the model. Genotype data were filtered for SNPs with minor allele frequency (MAF) > 0.1. Associations were tested using the ‘modelLINEAR’ setting, and cis-QTLs with FDR < 0.05 were retained for downstream analysis.

To identify potential disease-causal regulatory pathways linking genetic variants to chromatin accessibility and gene expression, we applied a relaxed effect significance threshold strategy. We first confirmed that gene - peak pairs linked by SNPs identified by MatrixeQTL have significantly higher correlations compared to gene - peak pairs not linked (**Fig. S12C**). Using this correlation enrichment as biological justification, we constructed a relaxed framework for identifying candidate SNP - chromatin - gene regulatory pathways. For each cell type, sex, and disease contrast, we filtered for SNPs that were significant in both cis-eQTL and cis-caQTL mapping (FDR < 0.05), and further required that the corresponding gene or peak showed differential expression or accessibility under disease-relevant conditions based on mashr estimated local false sign rate. Specifically, we retained triplets where either the gene or the peak exhibited strong differential signals (lfsr < 0.1) and the other exhibited suggestive but not definitive signals (lfsr < 0.5). This relaxed requirement aims to balance statistical stringency with biological sensitivity, allowing us to capture cases where one component may have a modest effect due to noise, variability, or indirect regulation. Gene–peak–SNP triplets passing these filters were considered candidate causal regulatory chains.

### Cells in droplet rate, cell multiplet rate, and cell hashing simulations

The Poisson distribution is well-suited for modeling cell encapsulation in droplets because it describes the probability of a given number of independent events occurring within a fixed volume when these events happen at random. However, using a Poisson model in isolation is insufficient for accurately predicting cell partitioning, as it does not account for cell loss and other experimental inefficiencies, such as droplets lacking barcoding beads.

To refine the Poisson model to predict real situations, we assumed that: (1) each 10X multiomics channel generates approximately 80,000 droplets^13,126^; (2) only 80% of these droplets contain barcoding beads^127^; and (3) around 10% of cells are lost due to experimental factors such as dead volume, pipetting loss, and sample handling (e.g., using only 70 µL of a 75 µL single-cell suspension master mix).

In this context, each cell independently enters a droplet: *X* ∼ *poisson*(*λ*), where *λ* represents the mean number of cells per droplet. *λ* is defined as *λ* = *I*/*N* · (1 − *α*), where *I* is the total number of input cells, *N* (80,000) is the total number of droplets, and *α* (0.1) is the fraction of cells lost due to experimental inefficiencies. To simulate droplets that lack barcoding beads, we randomly selected 80% of the droplets using a negative binomial model with a failure probability of 0.2. In each simulation iteration, we recorded the number of filled droplets. i.e., cell-filled droplets with (*X* > 0), singlet droplets (*X* = 1), and multiplet droplets (*X* > 1), with the multiplet rate calculated as the number of multiplet droplets divided by the number of filled droplets.

To evaluate the effect of multiplexing with hashing, cells were randomly assigned to unique hashing labels (e.g., 2, 4, 6, or 8) with equal probability. A droplet was classified as a detectable multiplet if it contained cells hashed with more than one unique hashing label. The proportion of detectable multiplets for each hashing condition was calculated as the number of detectable multiplets divided by the number of multiplets.

To assess the resilience of different multiplexing configurations to uneven hashing distributions, which increases the likelihood of homotypic multiplets, we introduced imbalance by systematically varying the probability of each hash’s occurrence, creating skewed distributions where certain hashes were overrepresented relative to others. This was achieved by defining a deviation parameter that modulated the uniformity of the probability vector assigned to hashes, with larger deviations corresponding to greater asymmetry in their abundances (**Fig. S3E**). When such a deviation parameter reaches 100%, one of the hashing barcodes will be completely absent.

Finally, we inferred the experimental mTEA-seq cell doublet rate based on the observed collision rate and the number of hashes used as:

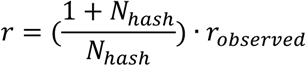

Where *r*_*observed*_ is the observed collision rate (calculated as the number of hash-detected doublet divided by the sum of doublet and singlet) and *N*_*hash*_ is the number of hashing labels used. The homogenous doublet rate was calculated as 1/*N*_*hash*_ times of the observed doublet rate.

## Supplementary Note

### Development of a multiplexed multiomic assay

Our approach utilizes antibody-based cellular hashing to allow for unambiguous multiplexing of samples, followed by FACS to deplete dead cells and granulocytes and allow for controlled pooling of samples, and finally multiomic processing of permeabilized cells on the 10X Genomics platform (**Fig. 1A**). In our initial experiments, we found that it could be difficult to unambiguously discern the neutrophils from the monocytes on the basis of CD15 alone during FACS sorting, and so we added CD16 to the staining cocktail to better distinguish monocytes and neutrophils (**Fig. S1A**). Additionally, we observed that co-amplifying cDNA and hashtag oligos (HTOs) in a single PCR reaction followed by separation using a double-size selection cleanup, per the original TEA-seq protocol, frequently resulted in non-specific byproducts in the HTO library, including aberrant fragment sizes and PCR bubble artifacts. This effect is likely attributable to the significant size disparity between HTO (∼200 bp) and cDNA (>600 bp), which leads to preferential amplification of the smaller HTO templates and consequent primer depletion. Although PCR bubble products are sequenceable, their presence introduces inaccuracies in fluorometric quantification due to the mixture of single- and double-stranded DNA structure. In contrast, conducting independent amplification reactions for cDNA and HTO markedly reduced the formation of such artifacts and simplified the protocol (**Fig. S2**).

As we intended to overload cells on the 10X microfluidic system (i.e. load more than the recommended number of cells and remove droplets with more than one hashing barcode detected *in silico*), we first sought to evaluate the collision rate as a function of cellular input using simulations. We modeled cell singlet and multiplet (i.e. more than one cell in a droplet) distributions for the microfluidic system with cell inputs ranging from 0 to 200,000 (**Fig. 1B**). Our simulations indicated that the total number of cell singlets peaked at a cell input of 94,949, which resulted in 23,959 singlets out of 42,263 recovered cell-containing droplets. However, at this level of overloading, 43.3% (18,304) of the recovered droplets sequenced would be multiplets, resulting in a large burden of unusable sequencing, as multiplets must be discarded when detected with hashing strategies. To account for this, we estimated the difference between singlets and multiplets and found that the yield of singlets over multiplets was maximized at an input of 48,484 cells, which would be expected to generate 20,584 singlet droplets and 6,509 multiplet droplets. In other words, accepting 14.1% fewer singlets results in elimination of 64.4% of unusable multiplets.

Another variable that can be adjusted is the number of hashing antibodies used in an approach like ours. While the number of hashing antibodies used will not affect the singlet and multiplet rates in our simulations, it does affect the likelihood of accurately detecting cell multiplets in an actual experiment without the use of additional doublet removal methods (i.e. DoubletFinder^128^ or demuxlet^129^). Because collisions can sometimes contain two or more cells with the same hashing barcode (called “homotypic” collision) not all collisions will be observable. Therefore, we sought to quantify the detectable multiplet fraction across different multiplexing configurations using simulations (**Fig. S3A**). Under idealized conditions with an even hashtag distribution and 48,484 cells as input (our previously identified optimum), detection of over 80% of multiplets requires the use of at least 6 hashing antibodies, while achieving more than 90% detection necessitates 10 hashing antibodies. However, these thresholds are predicated on uniform performance of hashes. In practice, technical variability (e.g., uneven antibody binding or differences in sample quality) can introduce imbalance in the hash distribution, resulting in increased homotypic multiplet rates (i.e., droplets containing multiple cells with identical hashing barcodes). To evaluate robustness against such an imbalance, we simulated skewed hash distributions by systematically modulating the occurrence probabilities (see details: Methods - Cells in droplet rate, cell multiplet rate, and cell hashing simulations). Our results indicate that employing 6 or more hashing antibodies maintained a multiplet detection efficiency of over 90% in skewed hash conditions (relative to a balanced hash representation), even when one sample was entirely lost (**Fig. S3A,E**).

Having determined reasonable parameters, we first sought to test our method with PBMCs from 4 individuals. Samples were processed and pooled equally and then we loaded 40,000 cells on the 10X microfluidic instrument. After sequencing, hashing tag oligo (HTO) signals were used for demultiplexing cells by sample source. Quantifying the relative abundance of HTOs across cells (**Fig. S3B**), we observed a clear bimodal distribution distinguishing cells with background rates of HTO reads from cells with clear positive signals for a specific HTO, suggesting antibody hashing specificity. Reassuringly, cells assigned to the 4 HTOs were approximately equally abundant. In total, we recovered 24,512 cell-containing droplets, of which 4,891 were classified as multiplets, 2,946 showed no detectable hashtag staining, and 16,675 were identified as singlets. Given that both mTEA-seq and TEA-seq were developed to enhance ATAC signal-to-noise ratios through the removal of neutrophils, we compared the ATAC quality metrics obtained from mTEA-seq with those reported for TEA-seq. We observed that the simulated cell multiplet rate closely matched the experimentally derived doublet rates from the mTEA-seq dataset (**Fig. S3C**). We also found our ATAC libraries were largely comparable to the original TEA-seq published data, with TEA-seq providing a higher “fraction of fragments in peaks” (FFiP) and mTEA-seq having a higher transcription starting site (TSS) enrichment scores (**Fig. S3D**).

With the 12,157 cells that passed further stringent RNA and ATAC quality filtering, we co-embedded the RNA and ATAC data in a unified Uniform Manifold Approximation and Projection (UMAP) plot (**Fig. 1C**). Despite shallow sequencing depth (a mean of 2,209 reads per cell for RNA and 3,734 read pairs per cell for ATAC), cells clearly separated into several major populations consistent with the expected cell types in PBMCs, including B cells, monocytes, and T lymphocytes. Cell clusters were annotated based on the expression pattern of selected cell type markers (**Fig. S3F**). The distribution of cells from each individual were relatively evenly distributed throughout clusters, suggesting minimal batch effects.

## Supplementary Figures

**Figure S1.**
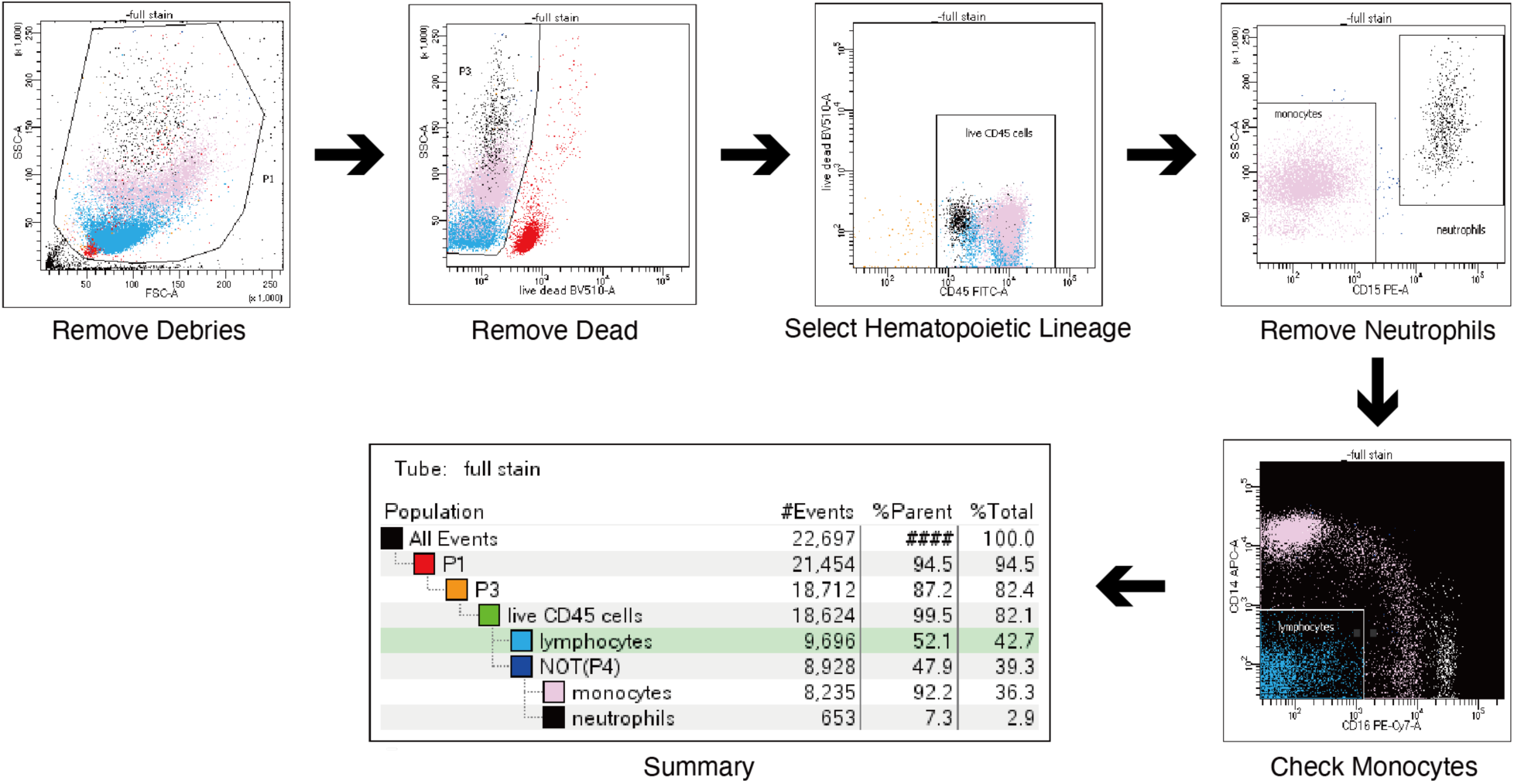
mTEA-seq PBMC sorting strategy. Sequential scatter plots illustrate the PBMC sorting workflow and neutrophil depletion strategy applied to mTEA-seq samples. From left to right: (1) debris exclusion based on SSC-A and FSC-A; (2) removal of dead cells using FVS-510 positivity; (3) enrichment for hematopoietic cells via CD45+ selection; (4) exclusion of neutrophils based on SSC-A and CD15 expression; (5) verification of monocyte retention by confirming CD14 and CD16 expression; (6) final gating summary with the proportion of cells retained at each step.

**Figure S2.**
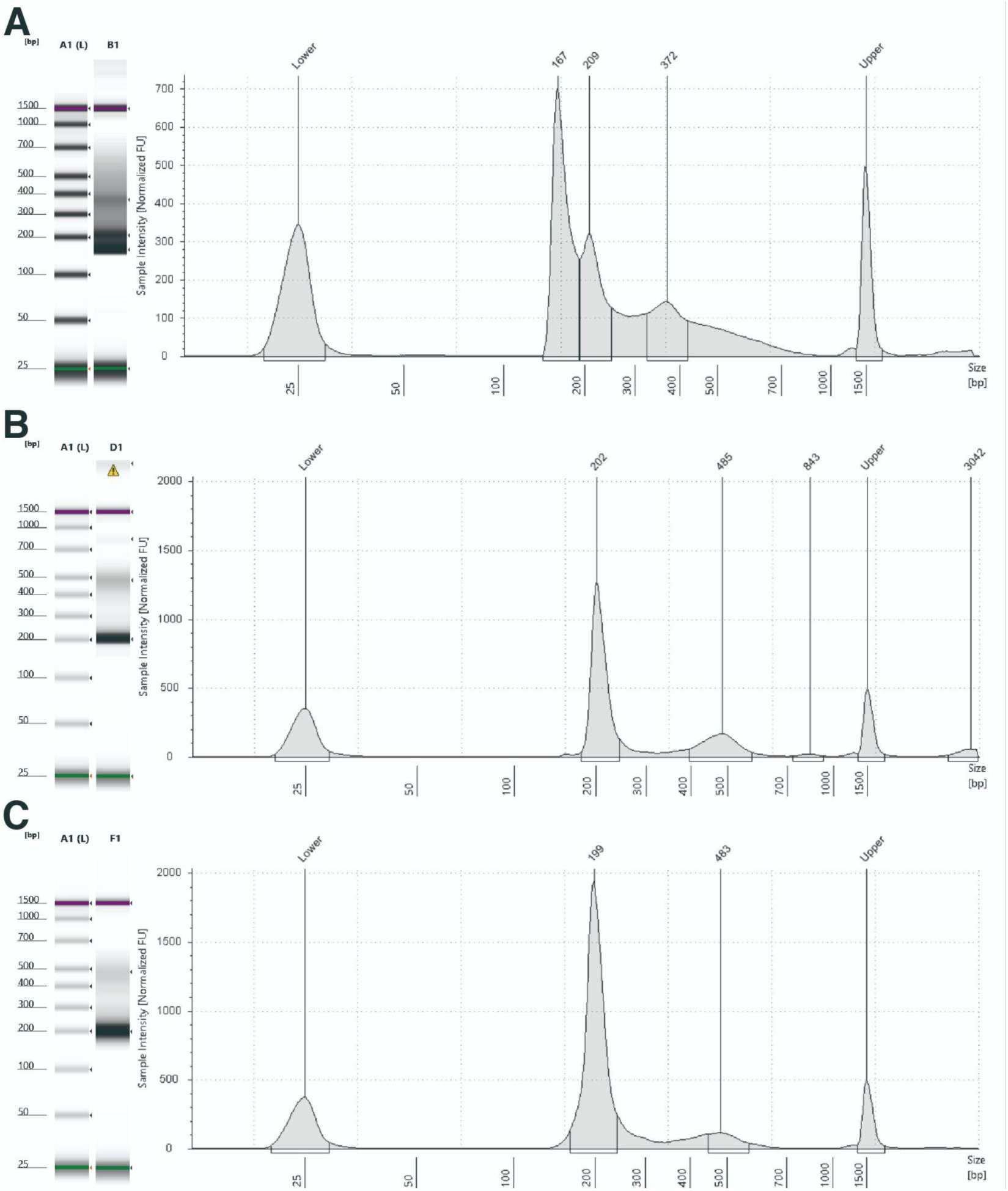
Direct amplification of hashing oligonucleotides mitigates the formation of PCR bubble byproducts. **(A)** TapeStation D1000 analysis of HTO libraries generated using the original TEA-seq protocol. The HTO library was co-amplified with cDNA and isolated with a 0.6X/2.0X double-sided size selection. The desired product appears at 209 bp. The size profile reveals non-specific PCR byproducts, including a prominent peak at 167 bp (likely primer-dimer) and a broader smear centered at 372 bp, indicative of PCR bubble artifacts. **(B)** TapeStation D1000 analysis of the HTO library in (A) following an additional 8 cycles of amplification with replenished generic P7 and P5 primers. The desired product appears at 202 bp, and PCR bubble byproducts are markedly reduced, with the residual artifact now observed at 485 bp. **(C)** TapeStation D1000 analysis of HTO libraries directly amplified from the pre-amplified mixed library (comprising ATAC, cDNA, and HTO components) using the improved mTEA-seq protocol. The expected product is detected at 199 bp, with minimal PCR bubble byproducts peaking at 483 bp.

**Figure S3.**
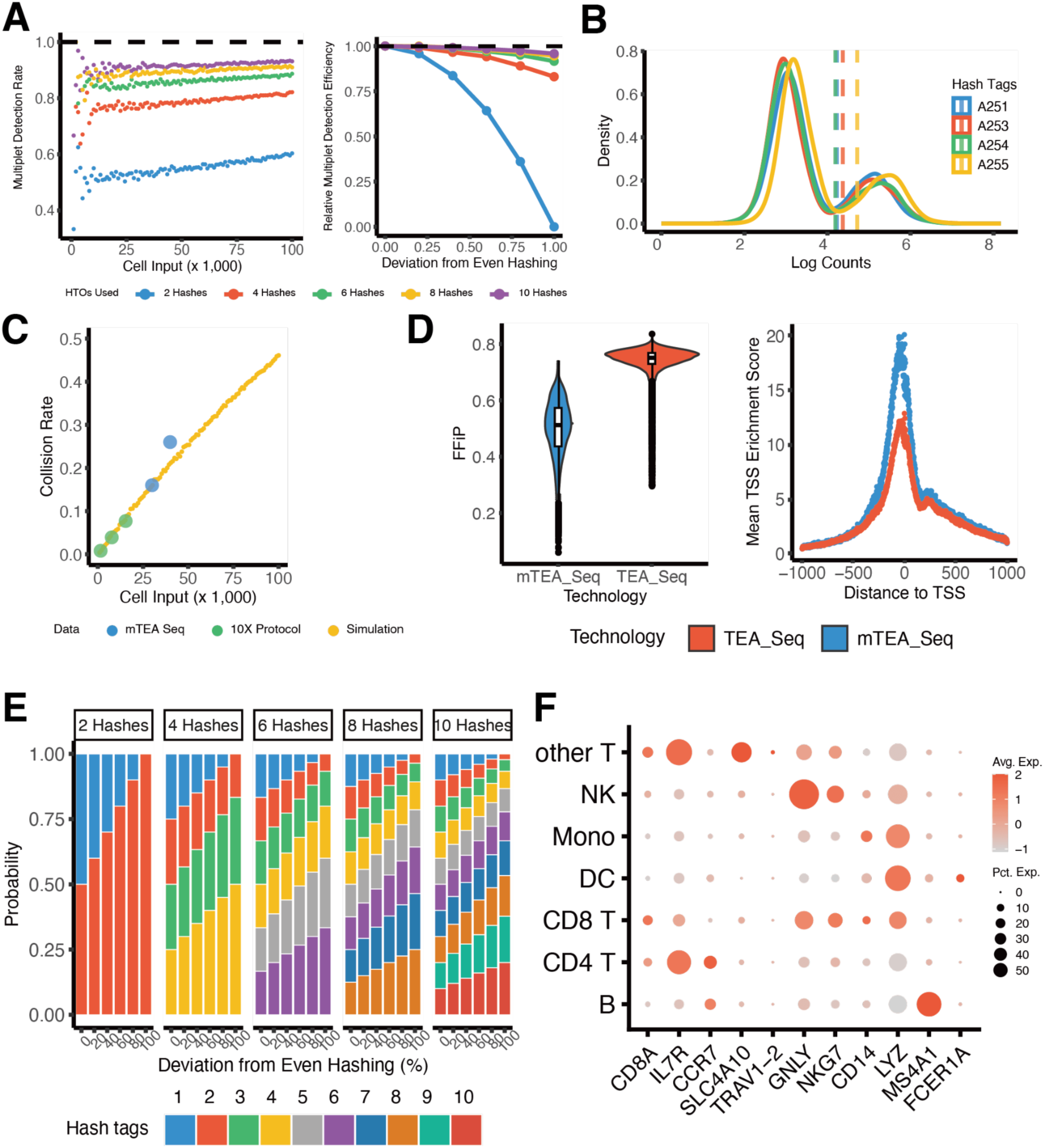
mTEA-seq simulations and quality controls. **(A)** (Left) scatter plot of cell input vs simulation-based multiplet detection rate (i.e., heterotypic collisions / total collisions) across various multiplexing configurations. Colors indicate the number of hashing barcodes, including 2 (blue),4 (red), 6 (green), 8 (yellow), and 10 (purple). (Right) Deviation from even hashing (i.e., fraction difference between the rarest and the most abundant hashing barcodes) vs the relative multiplet detection efficiency (i.e., fraction of multiplets detectable with perfectly balanced libraries that are still detectable under imbalance) across a range of states of imbalance, with the same color scheme as in (left) representing the corresponding number of hashing barcodes. **(B)** Density plots of number of HTO reads per cell for the four HTOs used on a logarithmic scale. Dashed lines mark the thresholds used to distinguish signal from background. **(C)** Scatter plot of cell input vs collision rate (i.e., fraction of droplets that are multiplets) for Poisson model simulation (yellow), alongside experimental collision rates obtained using mTEA-seq (blue) and the 10X protocol (green). **(D)** Violin plots (left) of the fraction of fragments in peaks (FFIP) for mTEA-seq (blue) and TEA-seq (red). Dot plots (right) ofTSS enrichment scores in a 2kb window around the start of genes. mTEA-seq (blue) and TEA-seq (red) are compared. TEA-Seq data was obtained from GEO accession GSM4949911. **(E)** Stacked bar plots of the distribution of sample proportions across different hash barcode configurations relevant to Fig. S3A, illustrating the progressively imbalanced representation of samples (see details in method section). **(F)** Dot plot displaying the expression profiles of canonical cell type marker genes across predicted cell types. Dot size represents the proportion of cells expressing each marker, while color intensity reflects the average normalized expression level.

**Figure S4.**
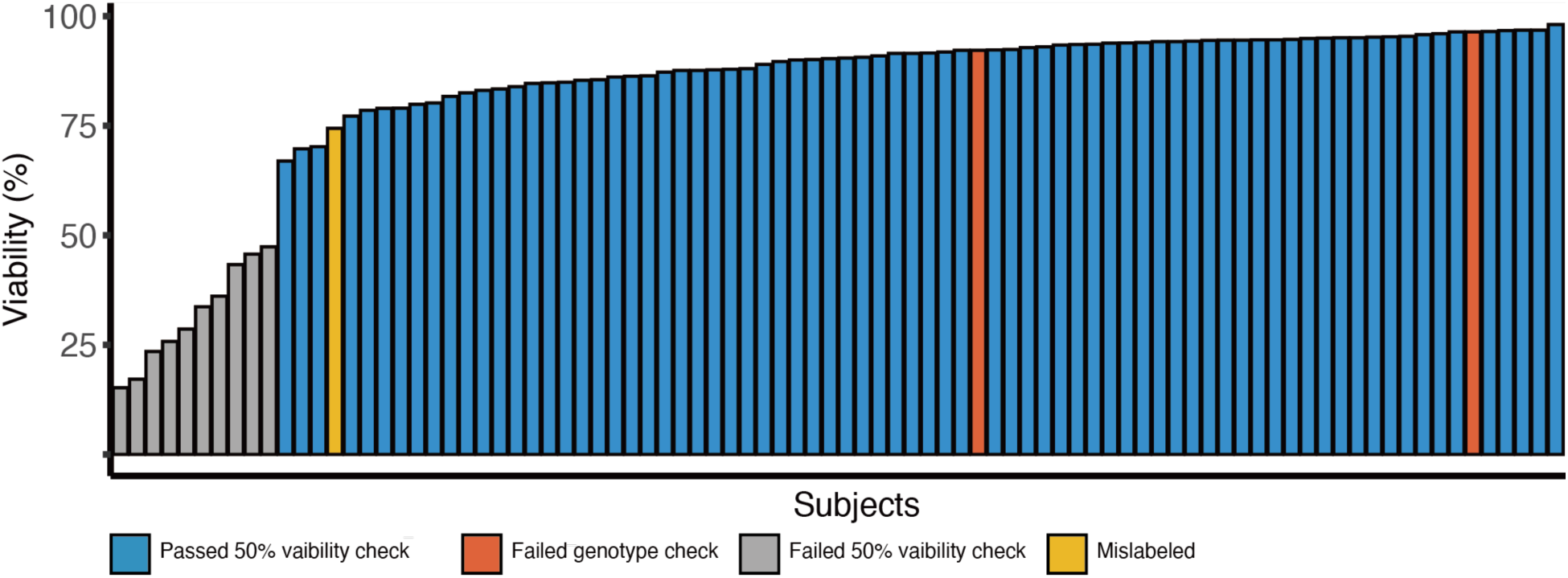
sample viability assessment in TCRS. Bar plots depict sample-level viability (determined by FVS-510 staining) across 89 PBMC samples, corresponding to 54 individuals from two batches. Samples with <50% viability were excluded: 6 from batch 1 (resulting in 33 retained) and 5 from batch 2 (resulting in 45 retained). One sample was too clumped to sort and so had no recorded viability.

**Figure S5.**
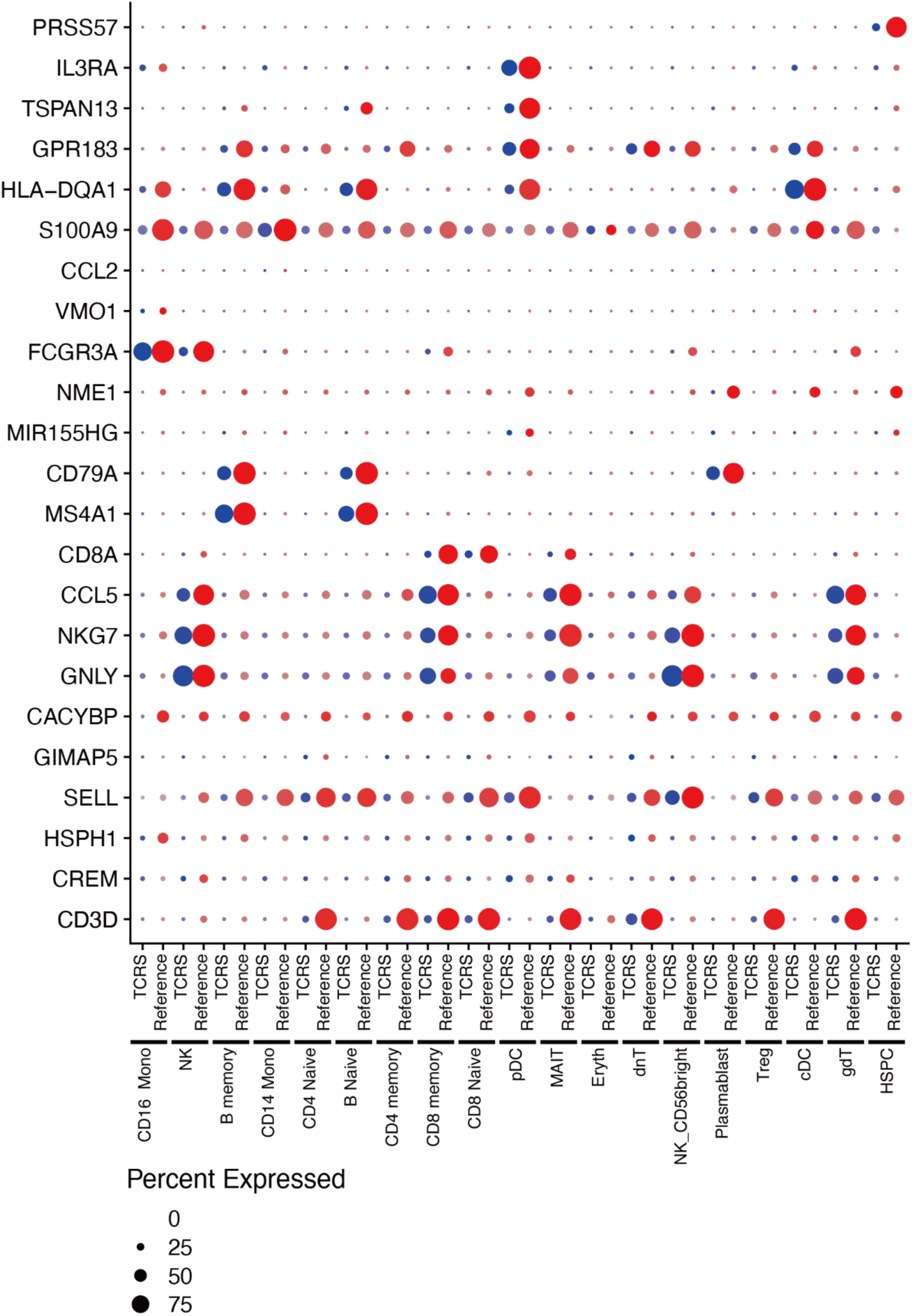
Cell type marker expression in TCRS PBMCs. Dot plots show expression of 23 marker genes across major cell types in TCRS single-cell PBMC samples (blue) and a reference PBMC dataset^107^ (red). Marker genes were curated based on established annotations provided in the Seurat vignettes^130^. Raw count matrices from both datasets were restricted to shared genes and overlapping cell-type annotations, then merged. The combined data were log-normalized, gene-scaled, and visualized using Seurat’s DotPlot function to compare expression levels and detection frequencies across matched cell populations.

**Figure S6.**
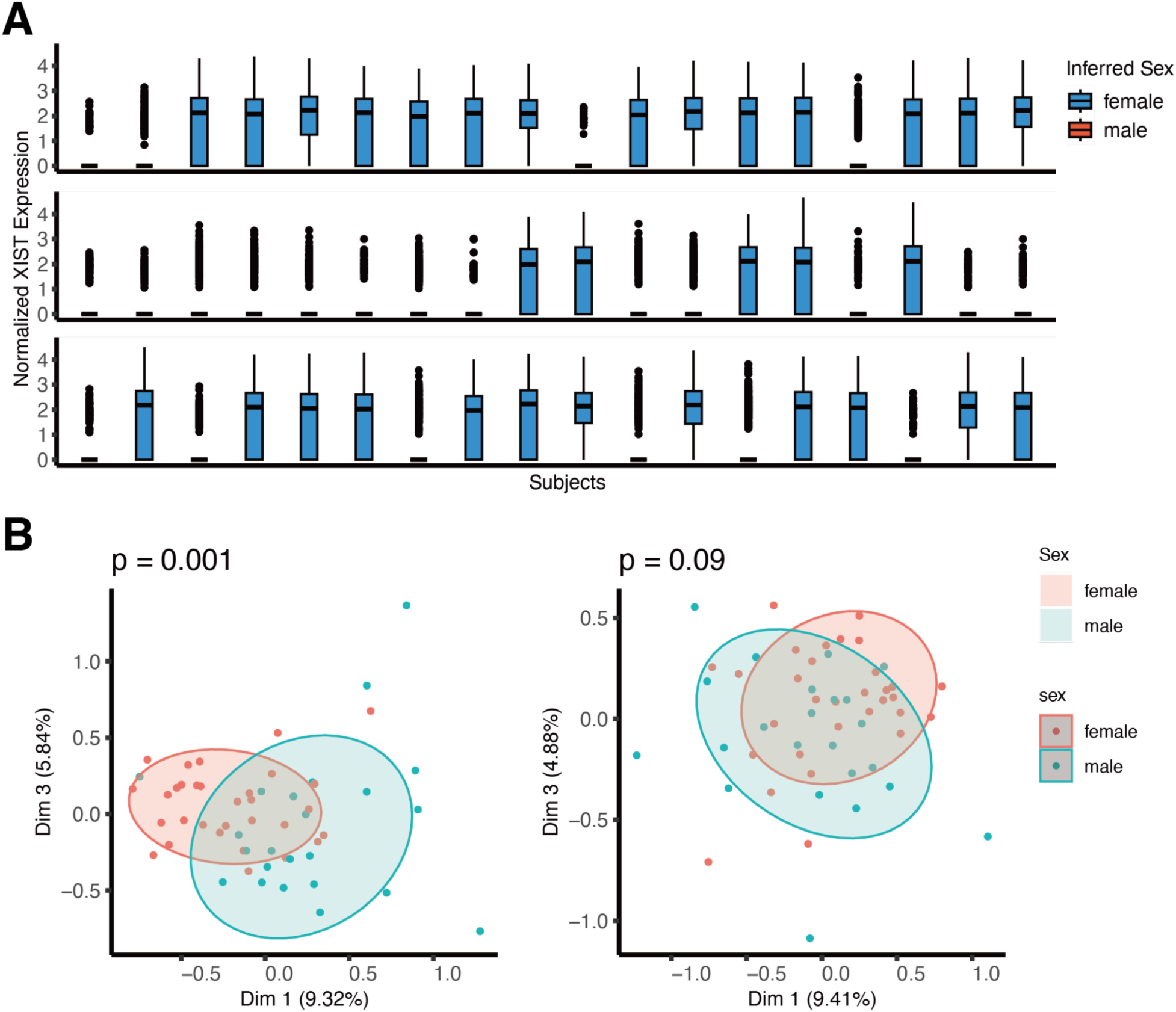
Pseudobulk RNA-seq PCA and pathway over-representation for male HN vs LN. **(A)** Boxplots of normalized XIST expression used to infer subject sex for the same 54 individuals. Blue indicates female, red indicates male. **(B)** PCA plots with Euclidean distance permutation testing show subject clustering by sex, with (left) or without (right) sex chromosome genes included. Green = male; red = female.

**Figure S7.**
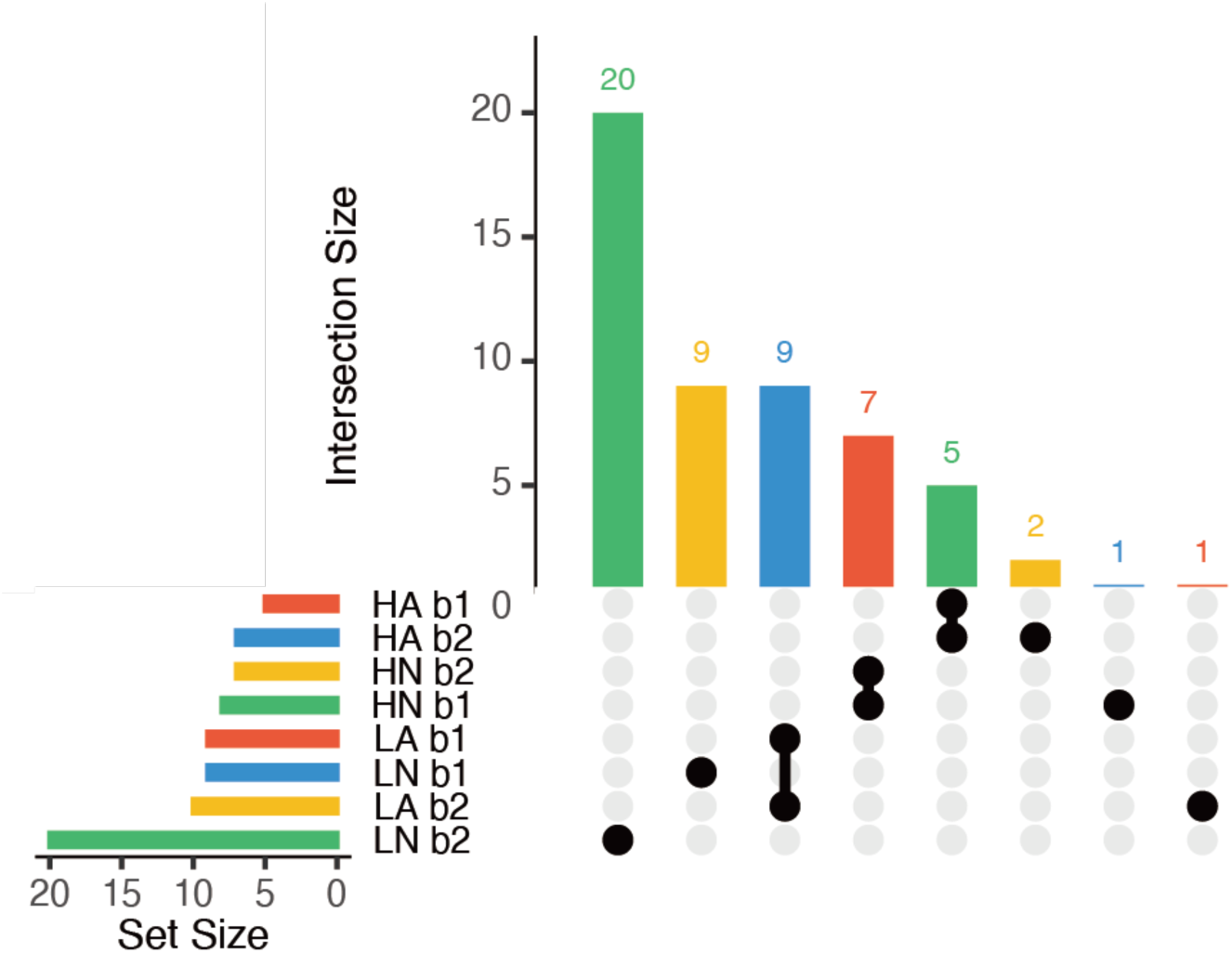
Subject overlap across batches. UpSet plot showing the distribution of shared and unique subjects across batch 1 (b1) and batch 2 (b2) for 75 high-quality samples (corresponding to 54 unique subjects). Colors indicate phenotype: green = LN; yellow = LA; blue = HN; red = HA.

**Figure S8.**
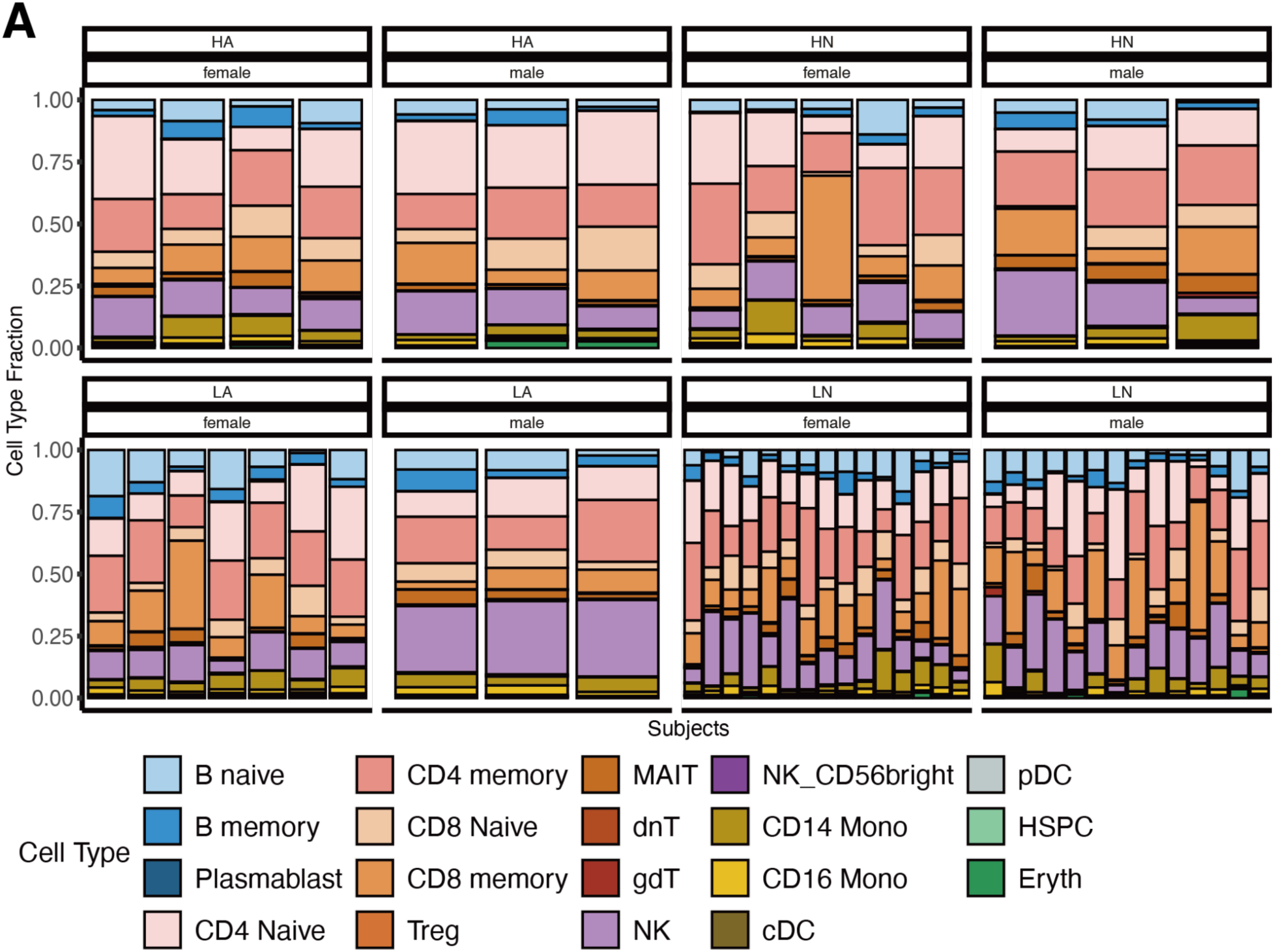
Cell type proportions across individuals split by sex and phenotypes. Stacked bar plot illustrating the proportion of annotated cell types for individuals across groups categorized by insulin and asthma status - low insulin non-asthmatic (LN), low insulin asthmatic (LA), high insulin non-asthmatic (HN), and high insulin asthmatic (HA) - further stratified by sex and phenotype. For individuals with repeated measurements across batches, the mean cell type fraction was calculated and used in the visual representation.

**Figure S9.**
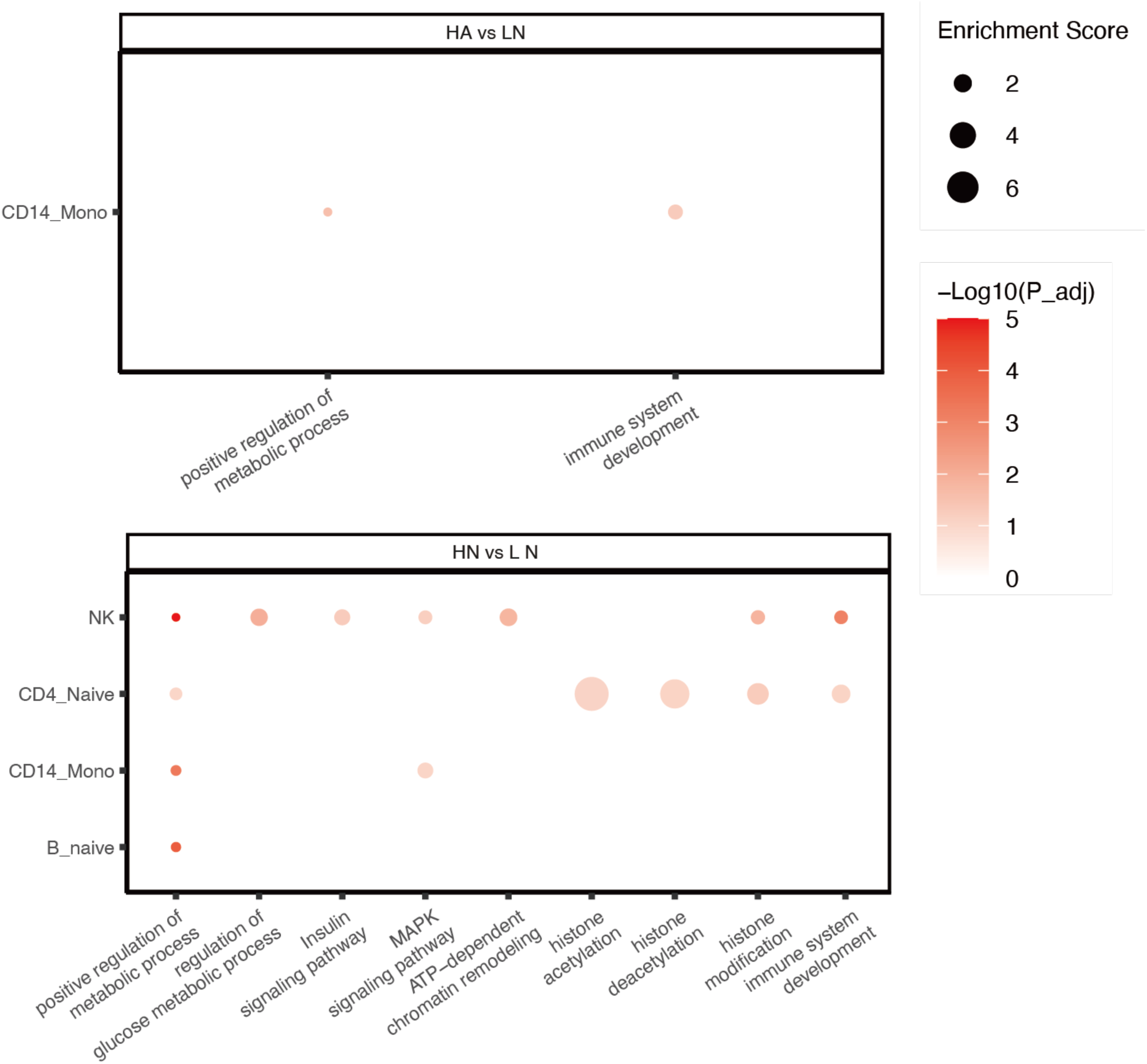
Pathway over-representation for male HN vs LN. Dot plots of gene ontology over-representation for cell types with over 100 differential accessible chromatin regions in the male HA vs LN and HN vs LN comparisons. The nearest gene for each DA peak was used for this analysis. Dot size reflects the proportion of genes overlapping with each pathway, and color indicates −log₁₀ adjusted p-values.

**Figure S10.**
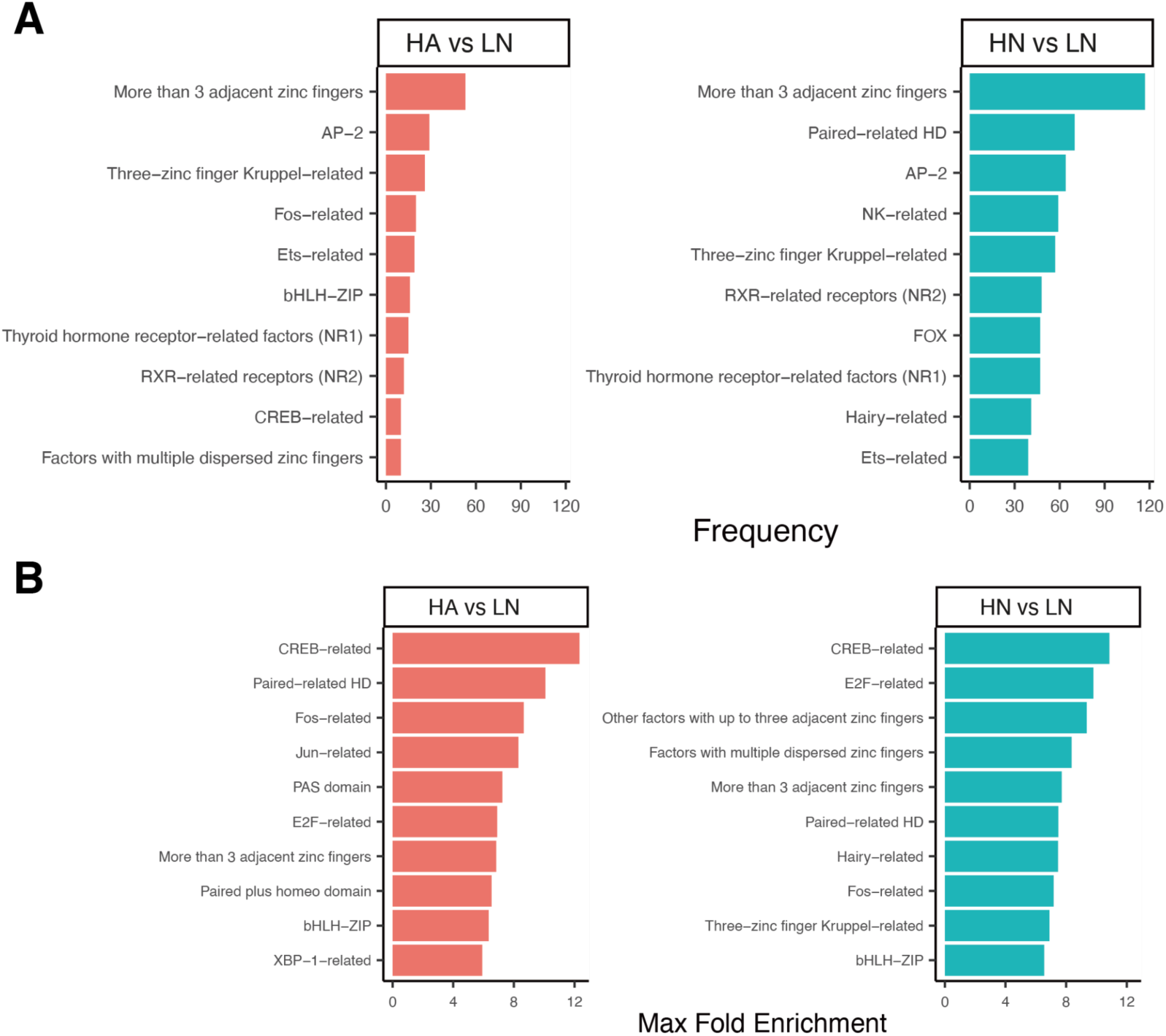
Top enriched transcription factor motif families identified from differential chromatin accessibility analyses. **(A)** Bar plots showing the ten most frequent motif families detected among active motifs (defined as enriched motifs corresponding to transcription factors expressed and exhibiting >1.5-fold expression difference between groups). Each facet represents an independent comparison: left, HA vs LN; right, HN vs LN. Bars indicate the number of distinct transcription factors per family. **(B)** Bar plots showing the top ten motif families ranked by the maximum fold enrichment of any motif within each family. Each facet again represents left, HA vs LN; right, HN vs LN.

**Figure S11.**
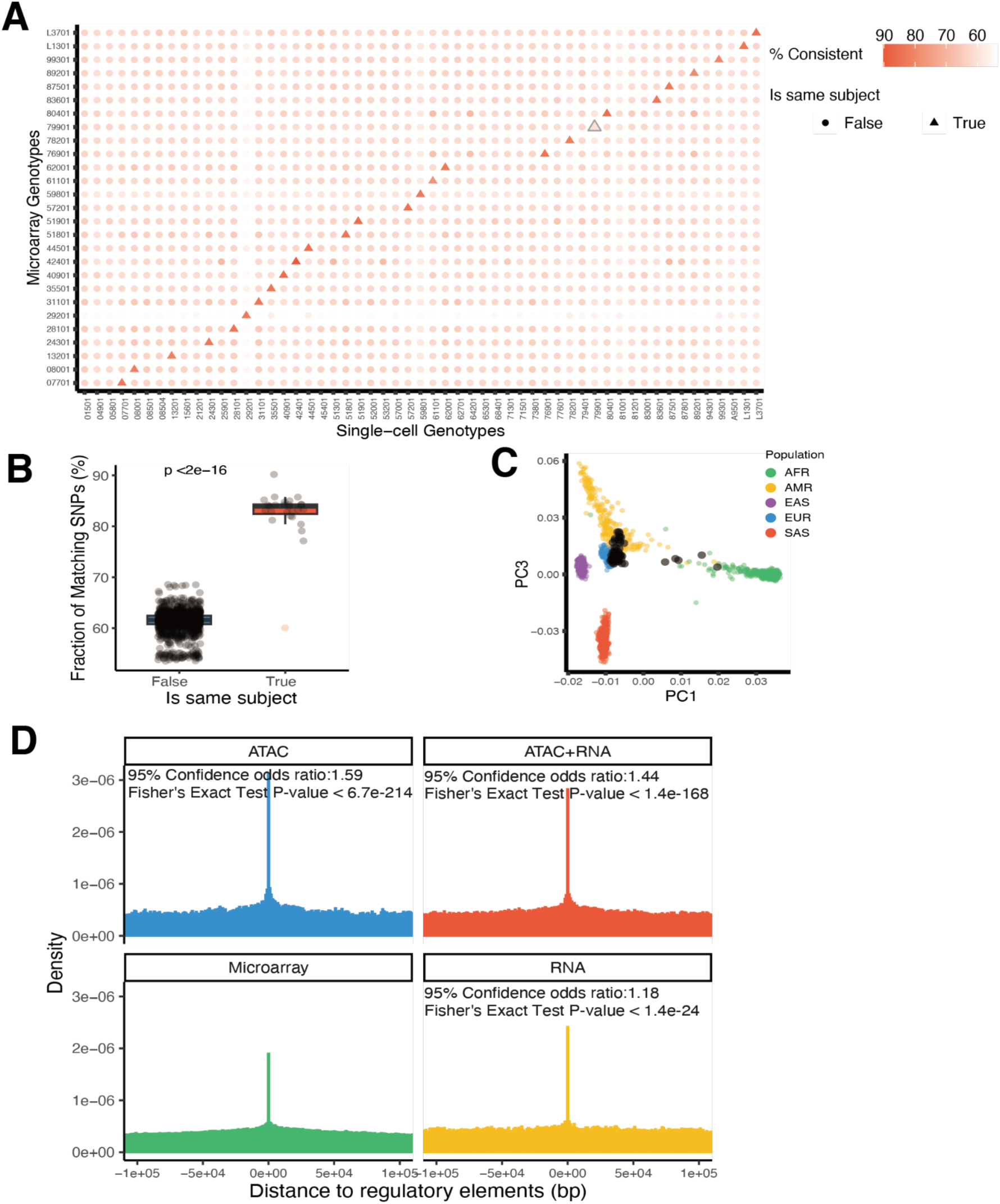
Concordance between single-cell and microarray-derived genotypes. **(A)** Pairwise comparisons of genotypes obtained from microarray data (27 subjects) and single-cell sequencing data (54 subjects). Data points are distinguished by shape, indicating whether the comparison was performed within the same individual (triangle) or across different individuals (circle). Color intensity represents the percentage of genotype concordance between platforms, ranging from low (white) to high (red). One subject exhibiting abnormally low self-consistency is outlined with a triangle. **(B)** Pairwise concordance rates between single-cell and microarray genotypes stratified by within-subject and across-subject comparisons. **(C)** PCA plots showing the genetic embedding of TCRS participants alongside individuals from the 1KGP reference populations. Genotypes for TCRS subjects were derived from single-cell multiomic data. Points are colored by population: African ancestry (AFR, green), Admixed American (AMR, yellow), East Asian (EAS, purple), European (EUR, blue), and South Asian (SAS, red). TCRS individuals are shown in black. **(D)** Density plots showing the distribution of distances between SNPs and reference functional regulatory elements regions. Genotypes were derived from the ATAC modality (blue), RNA modality (yellow), and combined ATAC+RNA (red) of the single-cell multiomic data, as well as from microarray-based SNP genotyping. Single-cell variants were identified using monopogen, while microarray SNPs underwent data cleanup (see Methods) and imputation.

**Figure S12.**
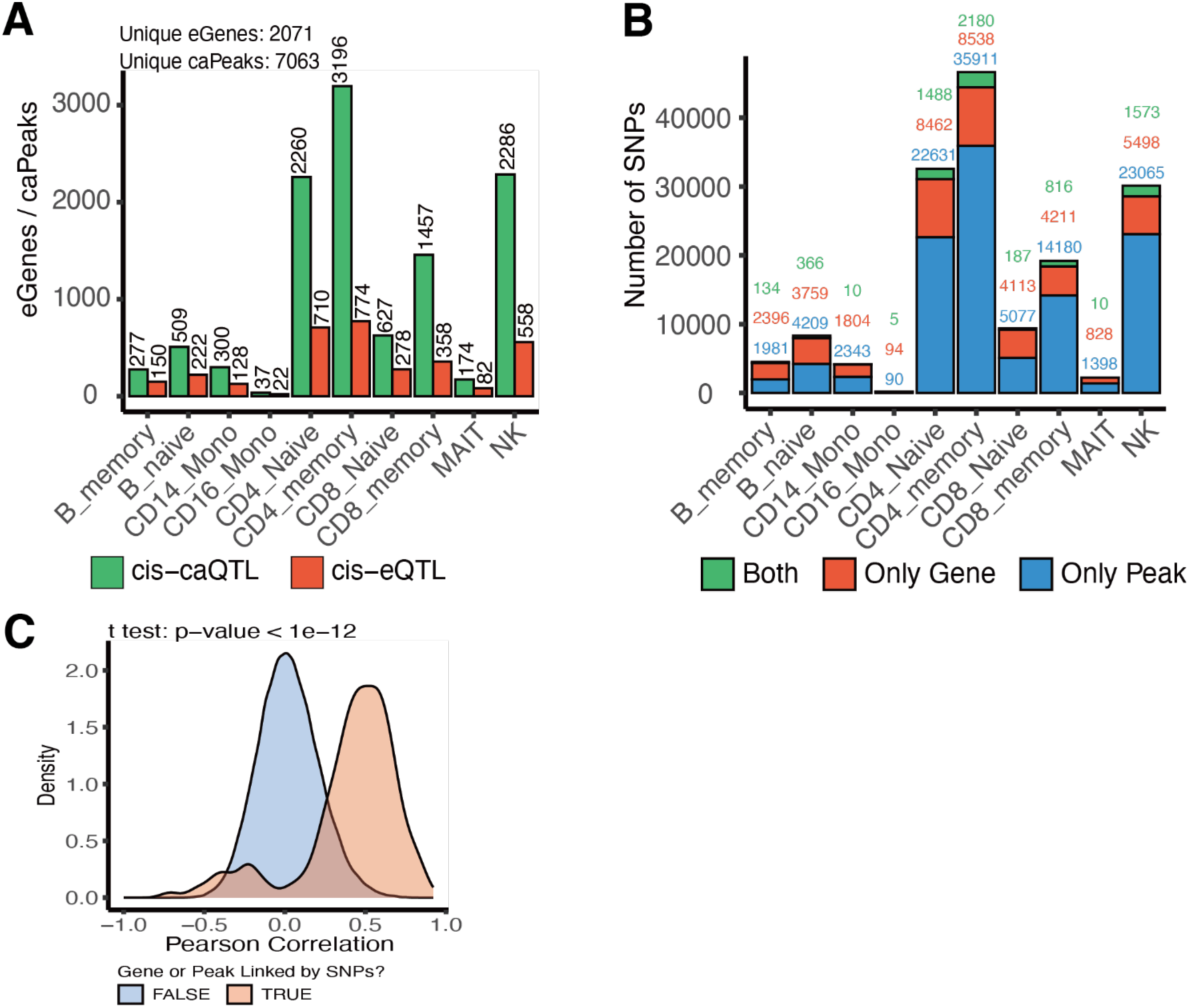
Cell type-specific cis-QTL discovery in TCRS single-cell multiomic data. **(A)** Bar plot showing the number of significant eGenes and caPeaks (FDR < 0.05) per cell type. Green = caQTL; red = eQTL. **(B)** Stacked bar plot summarizing the number of SNPs associated with genes, peaks, or both features in each cell type. Colors: red = gene only; blue = peak only; green = gene–peak pairs. **(C)** Density plot of the distribution of Pearson correlation coefficients for gene–peak pairs linked by shared cis-QTL SNPs (red), compared to random gene–peak pairs not associated through common SNPs (blue).

**Figure S13.**
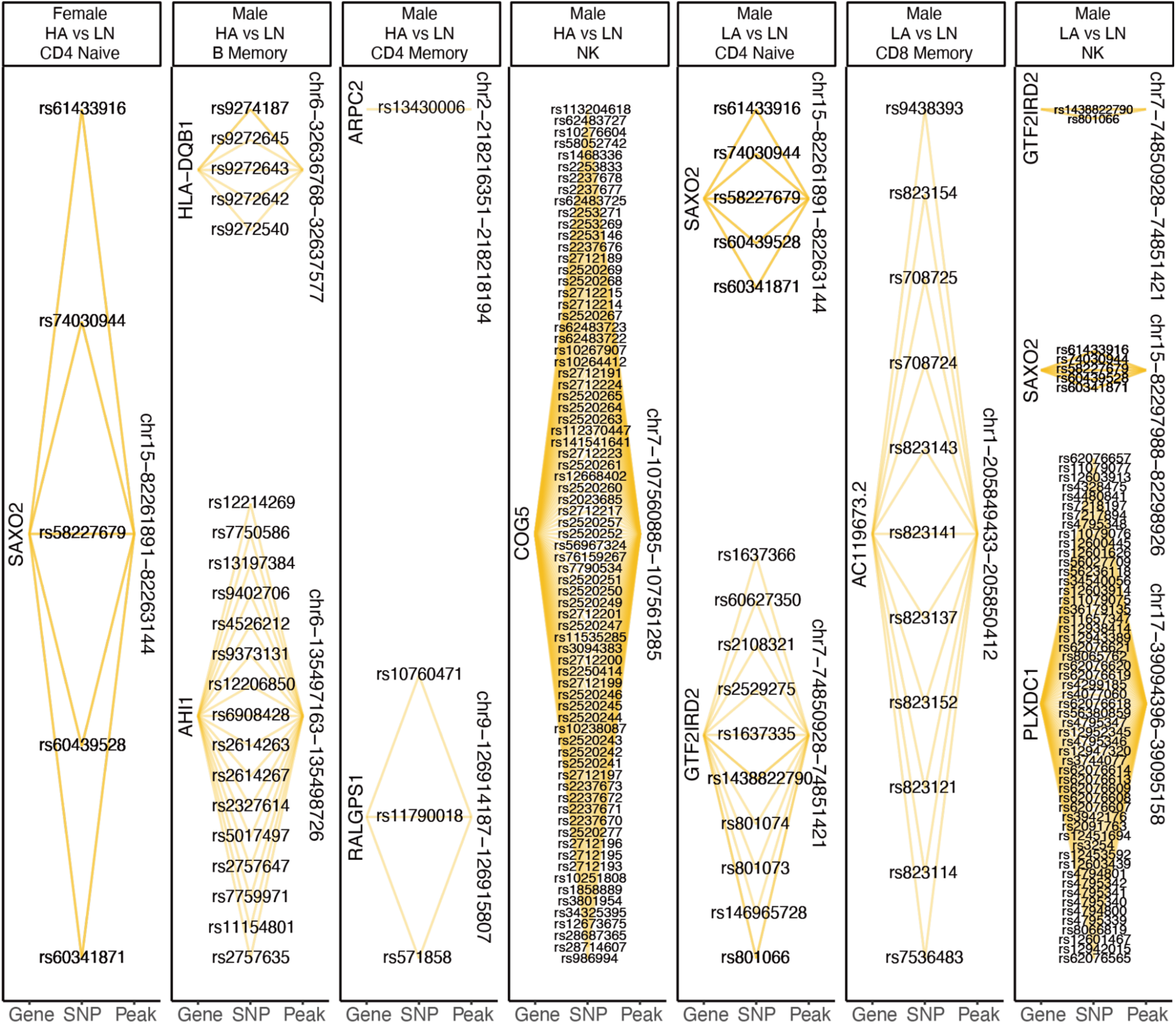
Identified gene - SNP - peak regulatory triplets. Summary of all gene–SNP–peak triplets that met the criteria of the two-threshold filtering strategy across all phenotype comparisons, stratified by sex. Each panel (from left to right) represents the following comparisons: HA vs LN in female CD4+ naïve T cells; HA vs LN in male memory B cells, male CD4+ memory T cells, and male NK cells; and LA vs LN in male CD4+ naïve T cells, male CD8+ memory T cells, and male NK cells.

**Figure S14.**
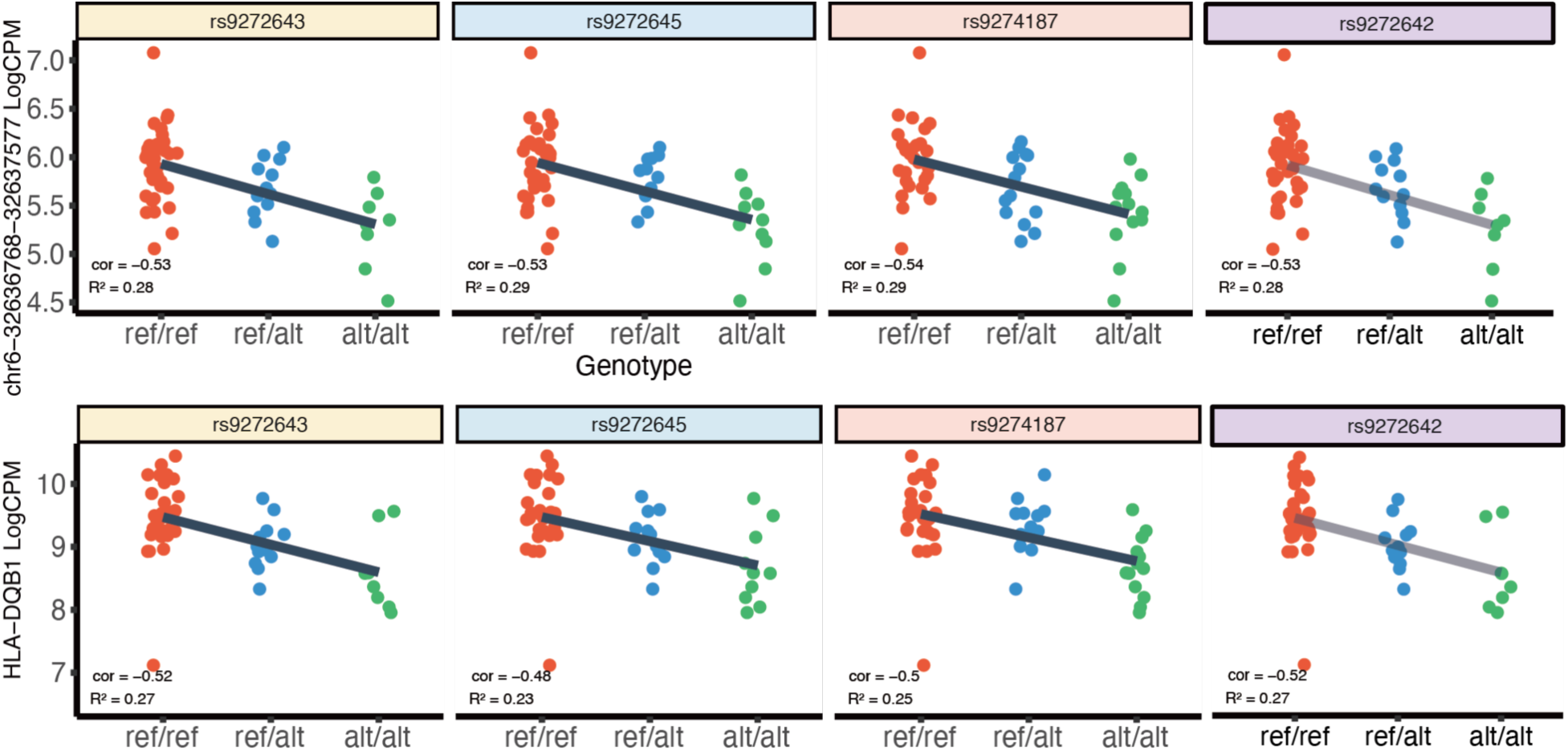
HLA Variants. Dot plots illustrating the relationship between genotype and molecular phenotypes for the four cis-acting SNPs (rs9272642, rs9272643, rs9272645, and rs9274187) not shown in Fig. 4B. The upper panels depict normalized expression levels of the HLA-DQB1 gene, while the lower panels show chromatin accessibility at the genomic locus chr6:32,636,768–32,637,577. Data points are stratified by genotype and color-coded as follows: reference homozygous (red), heterozygous (blue), and alternate homozygous (green).

**Figure S15.**
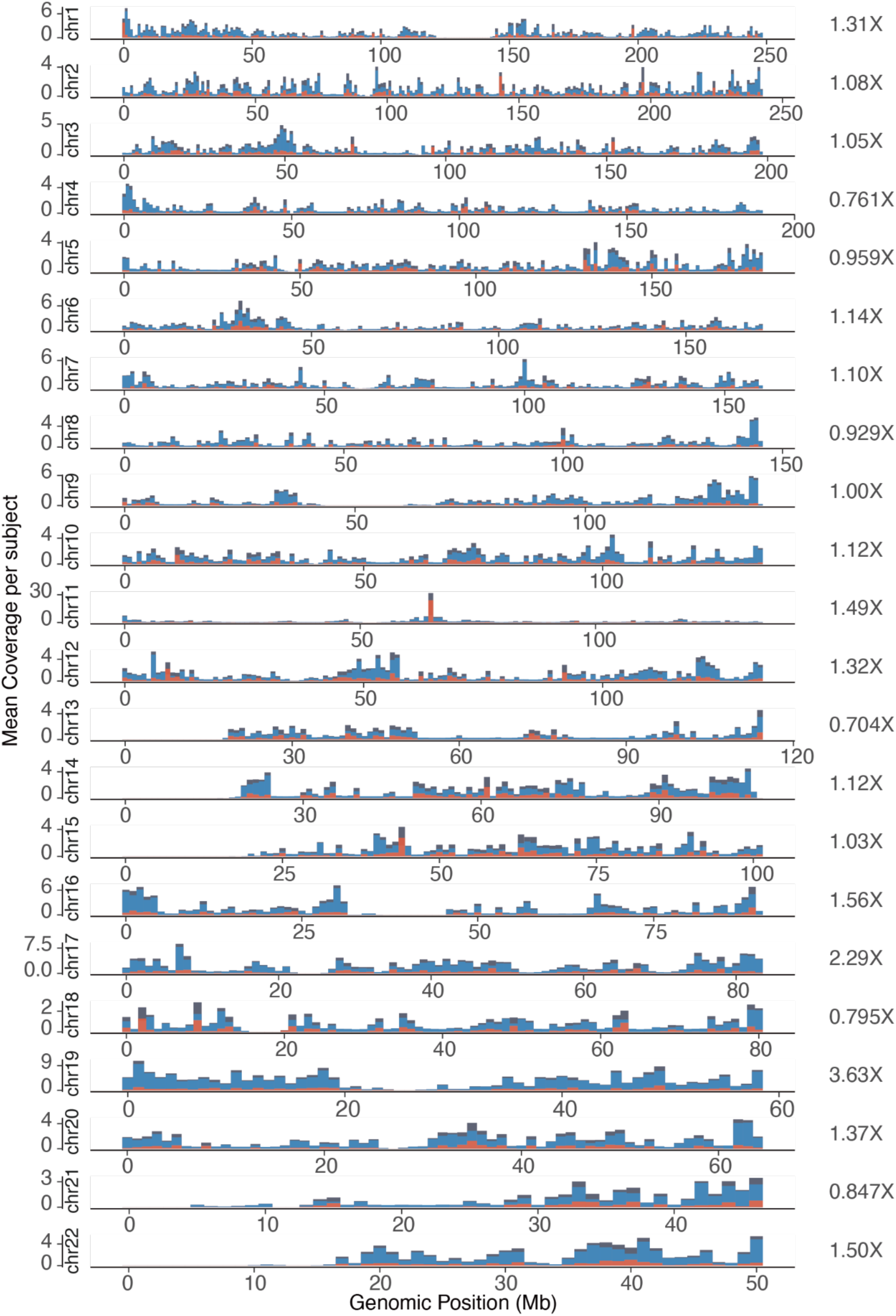
Genome-wide coverage of TCRS single-cell data. Bar plots showing aggregated chromosomal coverage across the 22 autosomes for RNA, ATAC, and merged single-cell datasets from all 54 subjects. PCR duplicates, including both optical and cell-level duplicates, were removed prior to coverage calculation. Data are color-coded as follows: red = RNA, blue = ATAC, and black = merged. The mean coverage per chromosome across all subjects is indicated to the right of each corresponding bar.

**Figure S16.**
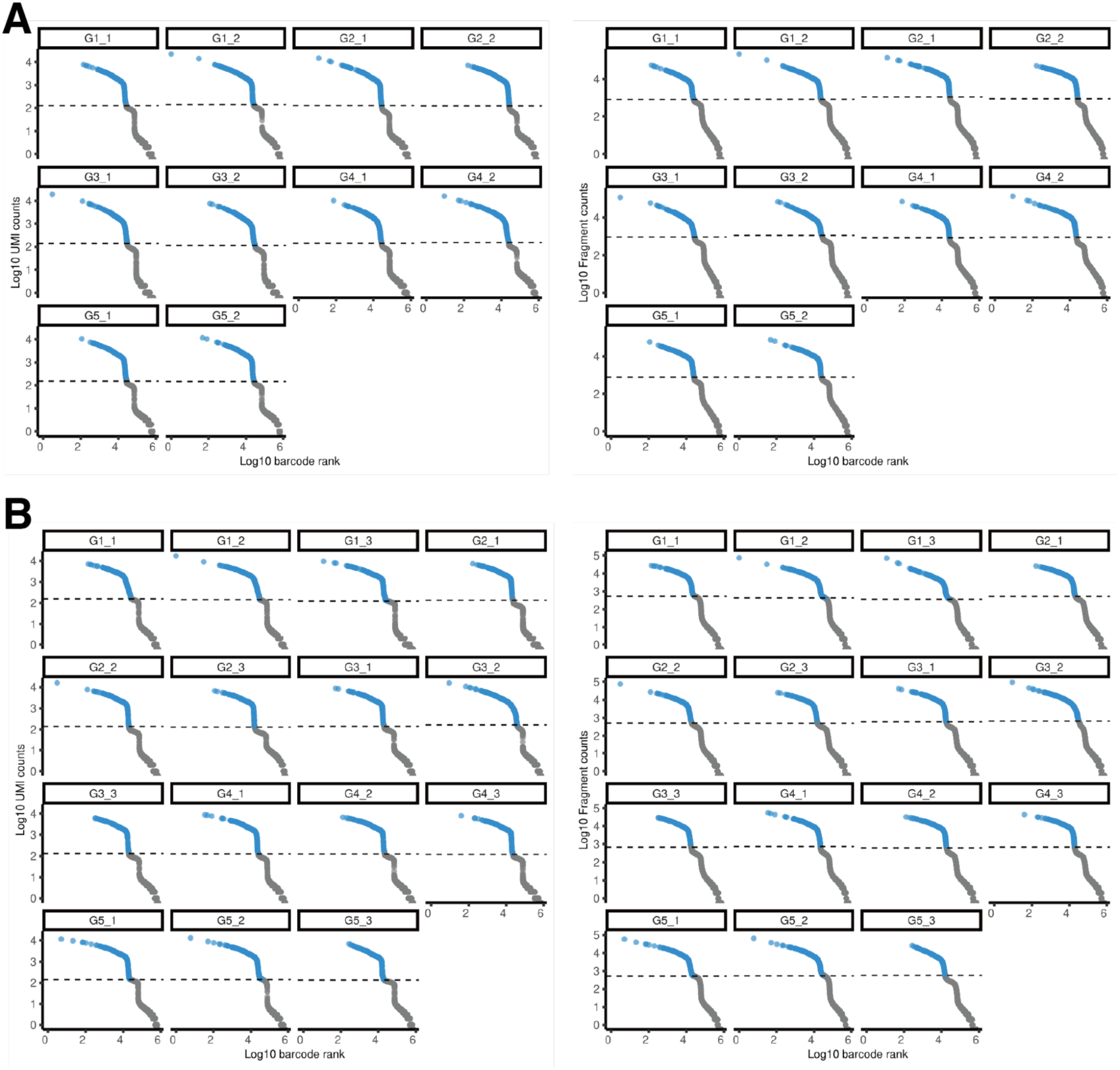
Knee plots for single-cell ATAC and RNA data in TCRS batches 1 and 2. **(A)** Knee plots display barcode-ranked log₁₀-transformed UMI counts (RNA, right) and fragment counts (ATAC, left) for batch 1. 10,000 barcodes were randomly selected for visualization, with cell (blue) and background barcodes (grey) clearly distinguished. Dashed lines indicate the threshold for distinguishing between cells and background barcodes, determined via k-means clustering per 10x Genomics channel.**(B)** Analogous plots for batch 2, with the same layout and processing parameters.

## Supplementary Table Legends

The Supplementary Tables are too large to include in the manuscript, and so are provided as separate files. Below, the columns of each table are described.

**Table S1. Sample metadata.**

A table contains metadata for each subject included in the study. It includes viability, batch, group assignments, and eligibility for downstream analyses. The table consists of 89 rows (samples) and 12 columns described below:

**subject_ID**: 5-digit subject identifier. This ID is consistent across batches and is used to match with clinical records.

**batch**: Batch in which the sample was processed (e.g., b1 and b2).

**live**: Percentage of live cells measured during FACS-based quality control. Expressed as a proportion (e.g., 94.5 means 94.5%).

**dead**: Percentage of dead cells, complement of the live column.

**subject_ID_unique**: Composite unique identifier for each subject-batch combination, formatted as “subject_ID + _ + batch” (e.g., 01501_b2).

**viability**: Indicates whether the sample passed FACS 50% viability QC. y: passed; n: failed.

**de_da_test**: Indicates if the sample was included in differential expression (DE) and differential accessibility (DA) testing. y: included; n: excluded.

**genotyping_qtl**: Indicates if the sample was included in single-cell based genotyping QTL mapping. y: included. n: not used.

**insulin**: Insulin status group of the subject. L: low insulin. H: high insulin.

**asthma**: Asthma diagnosis group. A: asthma. N: no asthma.

**ins_asma**: Combined phenotype label combining insulin and asthma status. Examples: low_yes, low_no, etc.

**comments**: Additional notes regarding sample inclusion or QC issues.

**sex**: Subject sex.

**Table S2. Significant mashr DE and DA analysis results.**

A table summarizes the results from mashr modeling applied to both RNA and ATAC assays. Each row represents a specific gene (RNA) or peak (ATAC) tested in a given phenotype comparison, stratified by sex and cell type. The table includes the following columns:

**gene**: For RNA-seq data, this refers to the gene symbol tested for differential expression. For ATAC-seq data, this refers to the genomic peak ID (e.g., chr:start-end) tested for differential accessibility.

**comparison**: The phenotype comparison performed. Examples: H_A vs L_N: High insulin with asthma vs Low insulin no asthma. H_N vs L_N: High insulin no asthma vs Low insulin no asthma. L_A vs L_N: Low insulin with asthma vs Low insulin no asthma.

**lfsr** (Local False Sign Rate): A Bayesian posterior error estimate computed by mashr that reflects the probability that the sign of the effect is incorrect.

**logfc**: The original log2 fold-change estimate from limma prior to mashr shrinkage. **true_bhat:** The mashr posterior mean estimate (shrinkage-corrected log2 fold-change). **cell_type:** The cell type where the differential expression or accessibility was tested.

**assay:** Indicates the type of molecular assay used. Values include: RNA: Single-cell RNA-seq data. ATAC: Single-cell ATAC-seq data.

**sex:** Indicates whether the contrast was tested in females or males.

**Table S3. Pathway overrepresentation results.**

A table summarizes pathway overrepresentation analysis results for gene sets derived from differentially expressed genes and differentially accessible peak nearby genes across various cell types, comparisons, and sexes.

**TermID**: The unique identifier for the pathway or biological process.

**Term**: The name of the pathway or GO term.

**Genes.in.Term**: The total number of genes associated with the term in the reference background.

**Target.Genes.in.Term**: The number of differentially expressed or accessible target genes that overlap with the term.

**Fraction.of.Targets.in.Term**: The fraction of target genes in the input set that are also in the term (Target.Genes.in.Term / Total.Target.Genes).

**Total.Target.Genes**: The total number of target genes tested for enrichment.

**Total.Genes**: The total number of genes in the background gene universe.

**Entrez.Gene.IDs**: The Entrez Gene IDs of overlapping genes in the term.

**Gene.Symbols**: The gene symbols corresponding to the overlapping genes in the term.

**p_value**: The unadjusted p-value from the enrichment test.

**adj_p_value**: The p-value adjusted for multiple testing, using Benjamini-Hochberg method.

**comparison**: The phenotype comparison from which the gene set was derived (e.g., "H_A vs L_N").

**cell_type**: The cell type in which the enrichment was tested.

**sex**: The sex of the donors in the comparison group (e.g., "male" or "female").

**assay:** Indicates the type of differential features used for pathway ORA test. Values include: “RNA” for differentially expressed genes or “ATAC” for differentially accessible peaks.

**Score**: Enrichment score defined as:

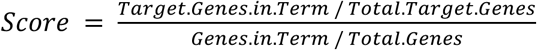

This ratio reflects the observed vs expected frequency of target genes in a given term.

**DataBase**: Indicates the source of the pathway annotation: either "GO" (for Gene Ontology Biological Process) or "KEGG" (for Kyoto Encyclopedia of Genes and Genomes pathway).

**Table S4. Significant QTL mapping results.**

A table summarizes the results of cis-QTL mapping performed across different immune cell types using either pseudobulk aggregated logCPM single-cell RNA or ATAC counts. Each row corresponds to a statistically tested SNP–target pair, where the target is either a gene (for RNA) or a peak (for ATAC). The columns include:

**rsid**: reference SNP ID annotated with NCBI dbSNP database reference (ftp://ftp.ncbi.nih.gov/snp/organisms/human_9606/VCF/All_20180418.vcf.gz).

**gene**: For RNA-based QTLs (cis-eQTL), this is the gene symbol of the transcript tested for expression association. For ATAC-based QTLs (cis-caQTL), this corresponds to the ID of the tested chromatin accessibility peak.

**beta**: The estimated effect size of the SNP on the molecular trait, derived from linear modeling, returned by MatrixeQTL.

**p.value**: The nominal p-value for the test of association between the SNP and the gene or peak, returned by MatrixeQTL.

**FDR**: The Benjamini-Hochberg adjusted p-value controlling for multiple testing across SNP–target pairs.

**cell_type**: The cell type in which the association was tested.

**assay**: Indicates the type of QTL being reported. Values include: “eqtl” for cis-eQTL; “caqtl” for cis-caQTL.

